# Tissue architecture dynamics underlying immune development and decline in the thymus

**DOI:** 10.1101/2025.06.28.662120

**Authors:** Sophia Liu, Mirco J. Friedrich, Ruth Raichur, Dennis Gong, Niklas Kehl, Qiyu Gong, Bingxian Chen, Karin Gustafsson, Larissa L. Ma, David T. Scadden, Rhiannon K. Macrae, Feng Zhang, Fei Chen

**Author notes:** Corresponding authors (S.L.), (M.J.F.), (F.C.). These authors contributed equally (Sophia Liu, Mirco J. Friedrich, Ruth Raichur).

## Abstract

The age-associated decline in adaptive immune function, widely observed in vertebrates, has been attributed to thymic involution. To gain insights into the structural and transcriptional changes underlying this phenomenon, we employed high-resolution spatial transcriptomics and T-cell receptor (TCR) sequencing in mice. By analyzing 21 thymus samples spanning mouse lifespan, we uncovered significant alterations in thymic organization, including disrupted T cell development and the emergence of B cell aggregates. We also observed age-related changes in cell-cell interactions, marked by increased antigen-presenting cell presence in thymic medullary regions and a shift from inflammatory to suppressive macrophages, fostering an immunosuppressive niche. Furthermore, aged thymus tissues exhibited an abundance of regions with reduced TCR diversity, accompanied by distinct changes to gene expression profiles. Our study establishes a valuable reference for understanding aging-related alterations in adaptive immunity, revealing mechanisms underlying age-induced immunological decline.

## INTRODUCTION

Immunological aging is a major contributor to age-related morbidity and mortality, impacting susceptibility to infectious disease and cancer, vaccine response, and autoimmunity^1–5^. Across nearly all vertebrates^6^, the immune system undergoes dramatic decline with age, leading to contracted adaptive immune repertoire diversity^7^, thus decreasing the threshold for immune escape and resulting in impaired immunity^4^. Age-related changes in immunity can be attributed to three sources: deteriorating hematopoietic progenitors^8^, declining organ/tissue structure^9^, and peripheral consolidation of T cell diversity following repeated antigen exposure^10,11^.

One of the most striking features of immune aging is thymic involution, the progressive shrinkage of the thymus, which is the primary site for T cell development and selection^12^. Thymic involution begins early in life — well before other signs of aging — resulting in a diminished output of naive T cells and a weakened immune repertoire. While the structural integrity of the thymus erodes over time^12–14^, leading to inefficient T cell neogenesis^15^, this process cannot be reversed by replenishing youthful hematopoietic precursors^16^, indicating that age-related changes in thymic architecture play a critical role in immune decline. Thymic involution not only affects immunity in aging individuals but also occurs in other contexts, including pregnancy^17,18^, acute infection^19–21^, and malnutrition^22^. Despite numerous single-cell studies characterizing longitudinal gene expression changes in the thymus^23–27^, these approaches have largely focused on dissociated cell states, offering limited insights into how tissue-level interactions shape T cell development. Although recent spatial transcriptomic studies have begun to reveal cellular interactions in the thymus, they have primarily focused on early development or disease states^28–30^. Constructing a high temporal and spatial resolution atlas across the trajectory of thymic involution would uncover age-related changes in cell interactions and identify potential targets to restore immune function.

Here, we aim to provide a comprehensive model of thymic and immune repertoire decline by integrating histological analysis, cellular composition, and T cell receptor (TCR) diversity. Our approach reveals age-related shifts in cell composition, diminished chemokine signaling, and the accumulation of thymic B cells. We also identify transcriptional changes in macrophages that disrupt their interactions with developing thymocytes, further compromising T cell development. Thus, by revealing how physiological aging-related changes couple thymus architecture to TCR repertoire diversity, we inform approaches to target and prevent immune aging.

## RESULTS

### Spatiotemporal profiling of changes in the thymus and peripheral blood throughout aging

To capture the dynamic changes to the thymus over thymic involution, we established a comprehensive spatial and single-cell atlas that not only maps TCRs and thymic architecture but also reveals critical shifts in cell composition, interactions, gene expression, and TCR organization throughout aging (**Fig. 1a**). Slide-TCR-seq^31^, a 10-µm-resolution spatial transcriptomics technique, uses 3 mm diameter circular arrays to capture cDNA from fresh-frozen, 10-µm-thick sections and enriches for TCR sequences. Using Slide-TCR-seq, we performed spatial transcriptomics and TCR sequencing on 47 spatial arrays from 21 mice aged from day 0 to week 90, characterizing 1,255,473 spatial locations; in parallel, we applied single-cell RNA and TCR sequencing (scRNA/TCR-seq) to profile 302,434 cells across eight time points in the thymus and peripheral blood, capturing systemic immunological changes (**Supplementary Table 1, Fig. 1a**). This approach enabled detailed analyses of cell composition and spatial localization, particularly within complex tissues where immune and stromal populations are differentially dissociated. Male mice were used to control for sex-based differences in thymic involution^32^, with at least two thymus sections sampled from a mouse per time point, either aggregating data across time points to achieve at least two mice per age group or performing trend-based statistical analyses using all time points.

**Figure 1.**
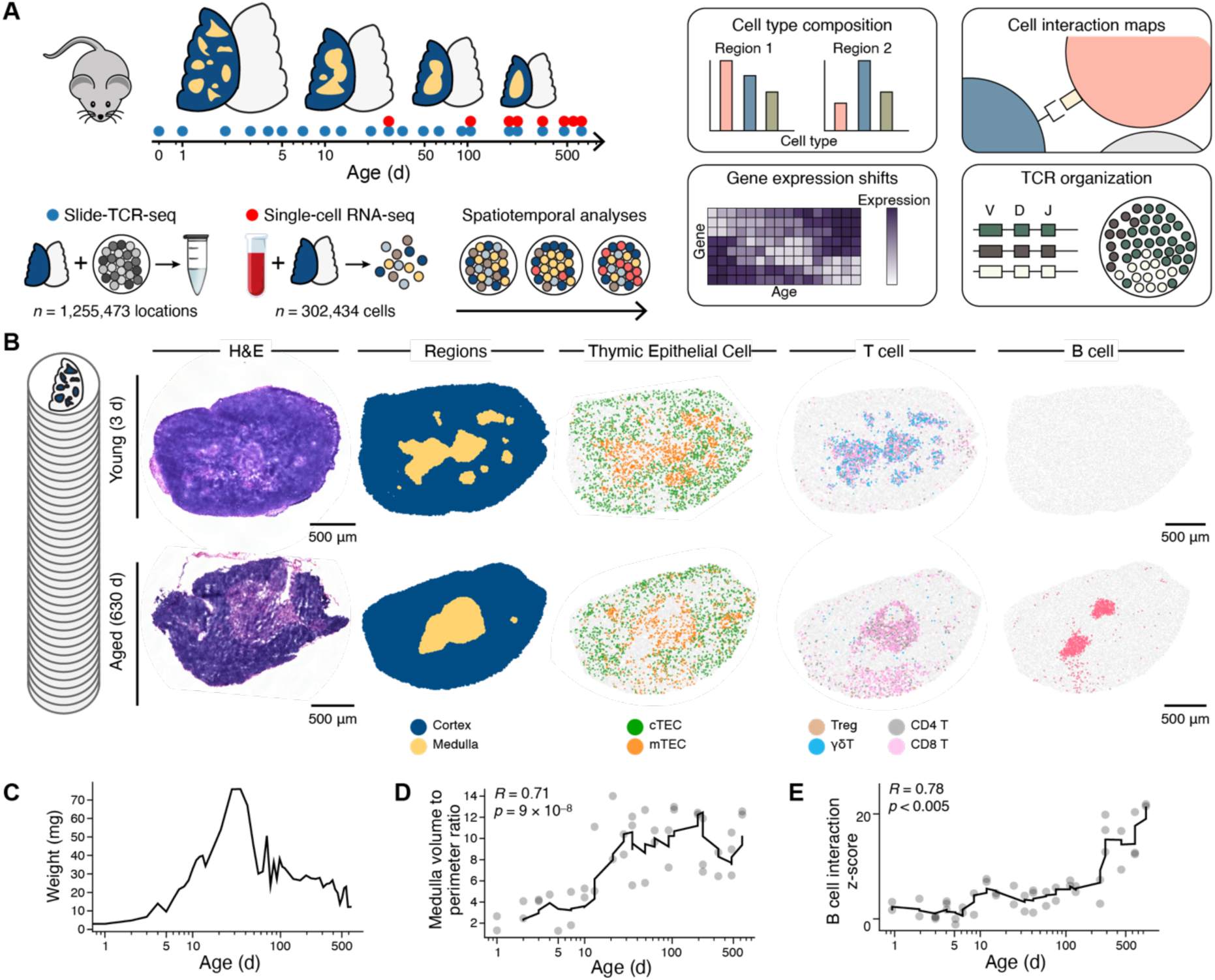
Spatiotemporal profiling of changes in the thymus and peripheral blood throughout aging. **a,** Changes to immune cell state and organ structure across the mouse lifespan identified using Slide-TCR-seq (n=47 spatial arrays over 21 mice/time points), and single-cell RNA-seq + V(D)J over 8 time points in the peripheral blood (n=175,839 cells) and thymus (n=126,595 cells), and overview of analyses. **b,** Slide-TCR-seq spatial arrays of representative young (3 d) and aged (630 d) mouse thymi. Left to right: **(i)** Hematoxylin and eosin stains of serial sections. **(ii)** Spatial arrays colored by cortex and medulla regions identified through unsupervised clustering. **(iii)** Spatial organization of cortical thymic epithelial cells (cTECs) and medullary thymic epithelial cells (mTECs). **(iv)** Changes to organization and types of T cells over age in the thymus. **(v)** Formation of B cell clusters over aging. **c,** Changes to thymic weight with age (n=47 mice). **d,** Changes in the consolidation of medullary regions in the thymus, as measured by surface area to perimeter ratio (n=41 spatial arrays over 20 mice). Rolling average line plotted, window = 5. Pearson’s r calculated using all time points. **e,** Significance of increased B cell spatial self-clustering over time determined via permutation testing (n=47 spatial arrays over 21 mice). Rolling average line plotted, window = 5.

We next used single-cell data^23,33^ to assign cell types to each spatial pixel using robust cell type decomposition (RCTD), a probabilistic model^23,34^. Unsupervised clustering identified cortical and medullary areas within the thymus, regions important for positive and negative selection of TCRs, and the transition of developing T cells from double-positive (DP) to single-positive (SP) states^35^. These regions were confirmed by hematoxylin and eosin-stained images, the localization of cortical and medullary thymic epithelial cells (cTEC and mTECs), and the presence of DP cells in the cortex and SP cells in the medulla (**Fig. 1b, Extended Data Fig. 1a**). Significant morphological changes in tissue architecture with age were observed: while thymus weight peaked at five weeks (**Fig. 1c**), medullary islands^36^ consolidated with age, as indicated by the ratio of surface area to perimeter of medullary regions (**Fig. 1b,d**, Pearson’s r = 0.71, p = 9.3 x 10^-8^). This structural loss coincided with shifts in T cell selection (decrease in DP to SP ratio with age, p < 0.005, by two-sample t test, **Extended Data Fig. 1b**), reduced Treg production (decrease in Treg to SP cell ratio with age, p < 0.005, by two-sample t test, **Extended Data Fig. 1c**), and aging-related cell state transitions in the medulla (**Extended Data Fig. 1d**, e). Notably, *Satb1+* expressing thymocytes in the medulla, which maintain the DP developmental state through chromatin architecture regulation^37^, increased with age, suggesting perturbed developmental dynamics (**Extended Data Fig. 1f**).

Our spatial data successfully recovered cell types typically difficult to detect in single-cell data, including stromal populations like mTECs. While our single-cell data contained 94.0% of total cells as thymocytes or T cells, enabling robust TCR clonotype detection and compartment tracing, 66.3% of spatial beads in our spatial data were thymocytes or T cells, demonstrating significant improvement in stromal cell capture (p = 1.9 x 10^-12^, by two-sample t test, **Extended Data Fig. 2a**). Eosinophils, granulated cells traditionally challenging to recover in single-cell sequencing due to rapid degradation, were also detected. Interestingly, eosinophils, which mediate thymic regeneration^38^, increased in the cortex with age but decreased in the medulla (**Extended Data Fig. 2b**), precluding eosinophil-mediated medullary repair over aging. We further observed the formation of B cell clusters over time (p < 0.005, Pearson’s r = 0.78, **Fig. 1b, e, Extended Data Fig. 2c**); building on mixed evidence^39,40^, we find that developing B cells, age-associated B cells, and mature B cells all increase with age in the thymus, supporting both peripheral recruitment and thymic B cell development (**Extended Data Fig. 2d**). Extending this analysis, we searched for the formation of irregular aggregates with age by measuring self-association and found that macrophages (p = 3.43 x 10^-5^, Pearson’s r = 0.62) and cDC2 cells (dendritic cell 2) (p = 0.00017, Pearson’s r = 0.57) similarly formed aggregates with age (**Extended Data Fig. 2e**).

### Age-associated shifts in cell composition and organization

Given the dramatic changes in the size and structure of the thymus with age, we anticipated shifts in both cell type proportions and cell-cell interactions. Across all time points, we observed that while the proportion of cells in the cortex that were cortical thymic epithelial cells (cTEC) remains relatively constant, the medullary thymic epithelial cell (mTEC) fraction in the medullary regions is displaced by antigen-presenting cells (APCs), broadly defined to include B, cDC1, cDC2, plasmacytoid dendritic cell (pDC), macrophage, and monocyte populations (**Fig. 2a, Extended Data Fig. 3a**, b). Dichotomizing time points into pre- and post-involution based on maximum thymus weight at 5 weeks, we found that double negative quiescent (DN(Q)) cells were predominantly in the cortex of pre-involution mice (p = 0.003, Cohen’s d = -1.6), whereas aged, post-involution mice had higher numbers of dendritic cell subsets (cDC1: p = 0.0086, Cohen’s d = 1.31; cDC2: p = 0.00056, Cohen’s d = 1.90; pDC: p = 0.005, Cohen’s d = 1.44) and NK cells (p = 0.00041, Cohen’s d = 1.45) (**Extended Data Fig. 3b**). In the medulla, we found that DN(P) cells (p = 0.00019, Cohen’s d = -2.21) and DP(P) cells (p = 0.0060, Cohen’s d = -1.46) were present significantly more in pre-involution mice, whereas post-involution mice had significantly more macrophages (p = 0.0019, Cohen’s d = 1.58) and cDC2 (p = 0.00042, Cohen’s d = 1.93) (**Extended Data Fig. 3b**).

**Figure 2.**
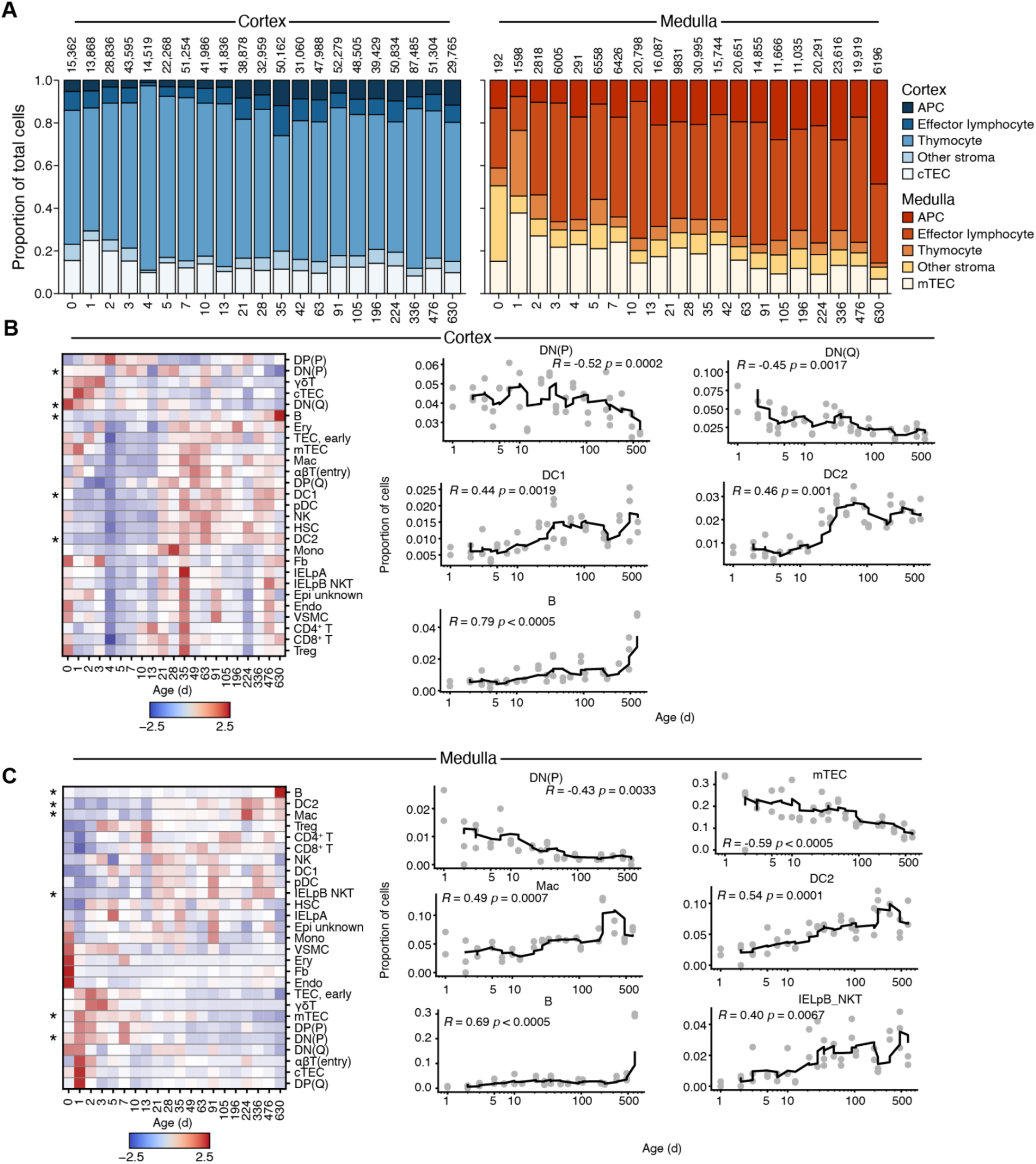
Age-related changes in cell composition. **a,** Cell type proportions over time in the cortex and medulla of antigen-presenting cells (APCs), effector lymphocytes, thymocytes, cortical and medullary thymic epithelial cells (cTEC, mTEC), and other stromal cells (n=21 mice). Numbers above the bar denote the total number of beads in each time point. **b-c,** Cell type proportions over time in the cortex (**b**) and medulla (**c**), separated by cell type (n=21 mice). Statistically significant shifts in cell type proportion after Benjamini-Hochberg correction are denoted by a * and visualized individually on the right. Rolling average line plotted, window = 5. Pearson’s r calculated using all time points.

By looking at individual cell types across the thymus and separately analyzing cortical and medullary regions, we revealed distinct age-associated shifts in cellular composition within each region (**Fig. 2b,c**, **Extended Data Fig. 3c**.) In the cortex, there was a notable decline in thymocytes (DN(P): p = 0.0002, R = -0.52; DN(Q): p = 0.0017, R = -0.45) and a corresponding increase in dendritic cell and B cell populations (DC1: p = 0.0019, R = 0.44; DC2: p = 0.001, R = 0.46; B cells: p < 0.0005, R = 0.79) (**Fig. 2b**). In the medulla, similar trends were observed, with a particularly strong reduction in mTECs (p < 0.0005, R = -0.59) (**Fig. 2c**). These findings underscore the substantial reorganization of thymic cellular composition with age, particularly the displacement of thymocytes and mTECs by immune cell populations, which may reflect decline in thymic output and immune tolerance as the thymus ages.

### Cell-cell interaction dynamics reveal changes in thymic architecture and macrophage states

To study how these differences in cell composition influence cell interactions over time, we next performed spatial bootstrapping (**Fig. 3a and Extended Data Fig. 3d**). In particular, in the cortex, we found that cTEC interactions with DN(P) cells decreased over time (p = 2.5 x 10^-3^, Pearson’s r = -0.54), whereas fibroblast interactions with CD4+ T cells increased with time (p = 8.9 x 10^-4^, Pearson’s r = 2.1 x 10^-3^), consistent with increased thymic fibrosis with age (**Extended Data Fig. 3e**). In the medulla, we observed an increase in mTEC interactions with DP(Q) cells over time (p = 3.8 x 10^-3^, Pearson’s r = 0.65), as well as an increase in interactions between mTEC and re-circulating αβT cells (p = 2.1 x 10^-3^, Pearson’s r = 0.72) (**Extended Data Fig. 3e**), possibly due to naive T cells in the periphery re-entering via leaky vessels^41^ into the aged thymus. To understand how T cell development may change with age, we developed a method for identifying changes in tissue structure and cell localization (**Extended Data Fig. 4**). For each time point, we identified the boundary by determining the alpha shape of the tissue^42^ (**Extended Data Fig. 4a**). We next measured the distance of the double-positive proliferating DP(P) and double-positive quiescent DP(Q) cells in the tissue to the boundary to build two distance distributions. Then, using the Kolmogorov-Smirnov test to determine if the two distance distributions for the cell types came from the same distribution (**Extended Data Fig. 4a**), we found that, consistent with visual observation (**Extended Data Fig. 4b**), at young, pre-involution time points, the two cell types occupied separate niches, reflecting the importance of spatial location and cell interactions in shaping the T cell developmental trajectory, which was significantly disrupted with age (p = 0.023 by permutation testing) (**Extended Data Fig. 4c**).

**Figure 3.**
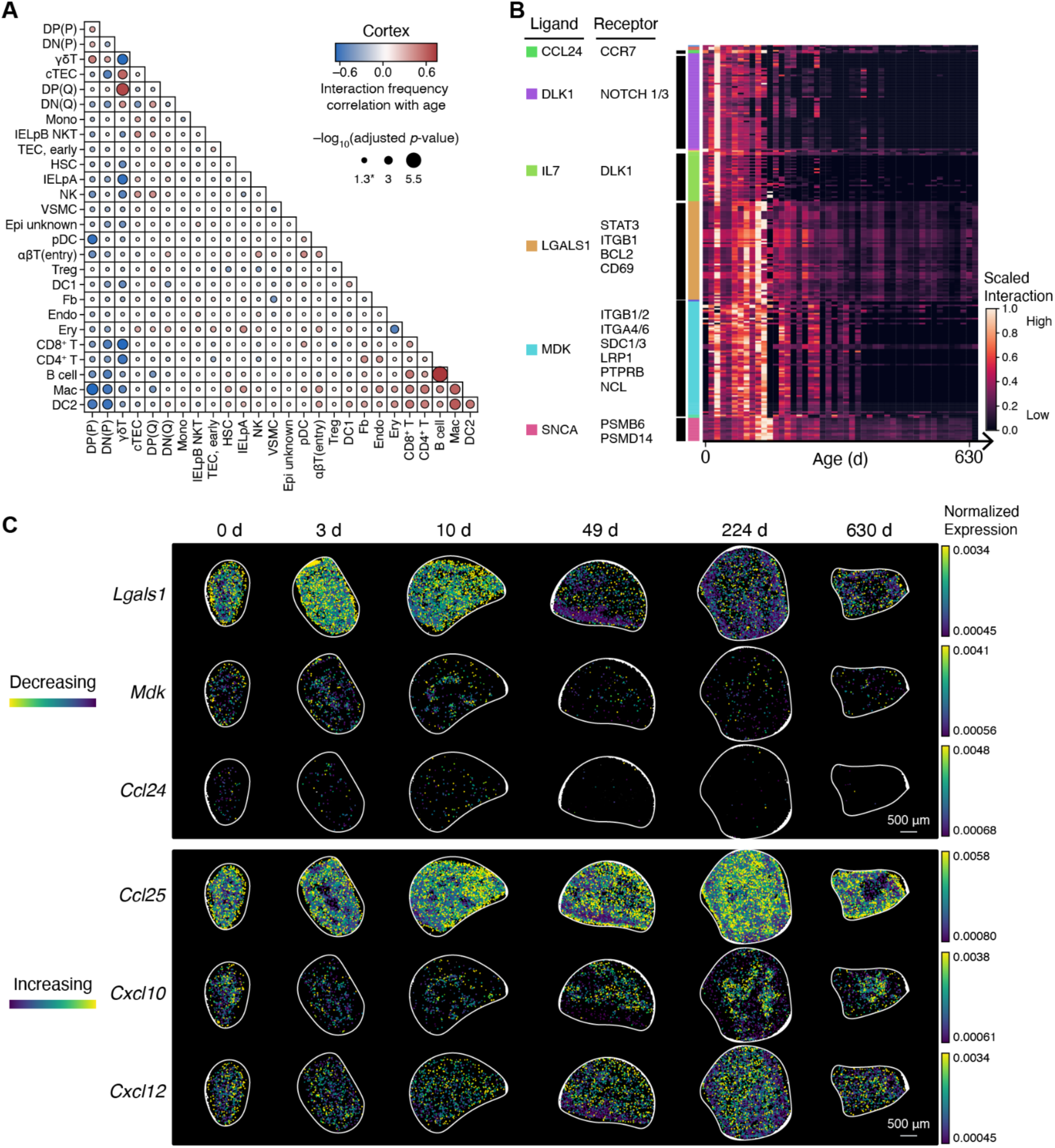
Age-related changes in cell interactions. **a,** Cell-cell interaction frequency correlation (Pearson’s r) with age in cortical regions of the thymus between all pairs of cell types (n=21 mice). Positive correlation values indicate increase in interaction with age. Circle size denotes adjusted p-value. **b,** Decreasing receptor-ligand interactions (normalized interaction strength, Methods) with age (n=21 mice). Each row is a unique cell-type-cell-type receptor-ligand pair and is annotated by common receptor-ligands. **c,** Visualizing selected receptor-ligand interactions that are decreasing and increasing with age. Color bar denotes normalized gene expression.

We next looked at receptor-ligand interactions to identify the specific cell interactions that may contribute to thymic involution. We used permutation testing, controlling for age-related changes in architecture and cell composition and creating a null model for cell interactions for each receptor-ligand pair via the Squidpy^43^ integration of CellPhoneDB^44^ and Omnipath^45^, to identify functional shifts in receptor-ligand interactions occurring over aging. Several receptor-ligand interactions were significant for decreasing over aging, most notably a decrease in interactions involving the expression of *Lgals1*^46^, *Mdk*^47^, and *Ccl24*^48^, previously associated with promoting a suppressive macrophage state (**Fig. 3b and Extended Data Fig. 5a**). We also observed increased *Ccl25, Cxcl10, Cxcl12,* and *Vcam1* expression with age (**Extended Data Fig. 5b**,c). *Vcam1* is important in guiding the homing of hematopoietic stem and progenitor cells, suggesting that the thymus may increase its recruitment of progenitors with age^49^. Visualized in the tissue context by plotting the normalized gene expression of the ligands, we observed the changes in the expression of these ligands in the spatial data (**Fig. 3c**).

### Decline in thymic TCR diversity with age

A key function of the thymus is to facilitate the generation and selection of TCR diversity^50–53^. To identify systemic effects of aging on T cell development, we first analyzed the scRNA-seq data and performed batch correction using Harmony^54^ (**Extended Data Fig. 6**). While cell states were generally shared across time points (**Extended Data Fig. 6e**,f), significant shifts in the proportions of certain cell types were observed. In peripheral blood, the proportion of CD4+ T cells decreased (p = 0.00019, Pearson’s r = -0.96), while NKT cells (p = 0.02804, Pearson’s r = 0.76) and CD8+ T cells (p = 0.0002, Pearson’s r = 0.96) increased (**Extended Data Fig. 6e**,f). These changes were mirrored in the thymus, where whole-tissue dissociation revealed a slight shift from CD4+ to CD8+ T cells, consistent with previous findings (**Extended Data Fig. 6g**)^41,55^.

To further investigate TCR diversity, we downsampled each time point to 7,500 T cells in the blood scRNA-seq data and observed a reduction in T cell clonotype diversity in the peripheral blood (**Fig. 4a and Extended Data Fig. 7a**). This reduction was primarily driven by the clonal expansion of cytotoxic CD8+ T cells (**Extended Data Fig. 7b**). Analysis of our spatial thymus data, downsampled to 200 TCRs per time point, similarly revealed that TCR diversity decreased with age (p < 0.005, by two-sample t test, **Fig. 4b)**, and we hypothesized that spatial data would provide more insight into local differences in T cell diversification in the thymus.

**Figure 4.**
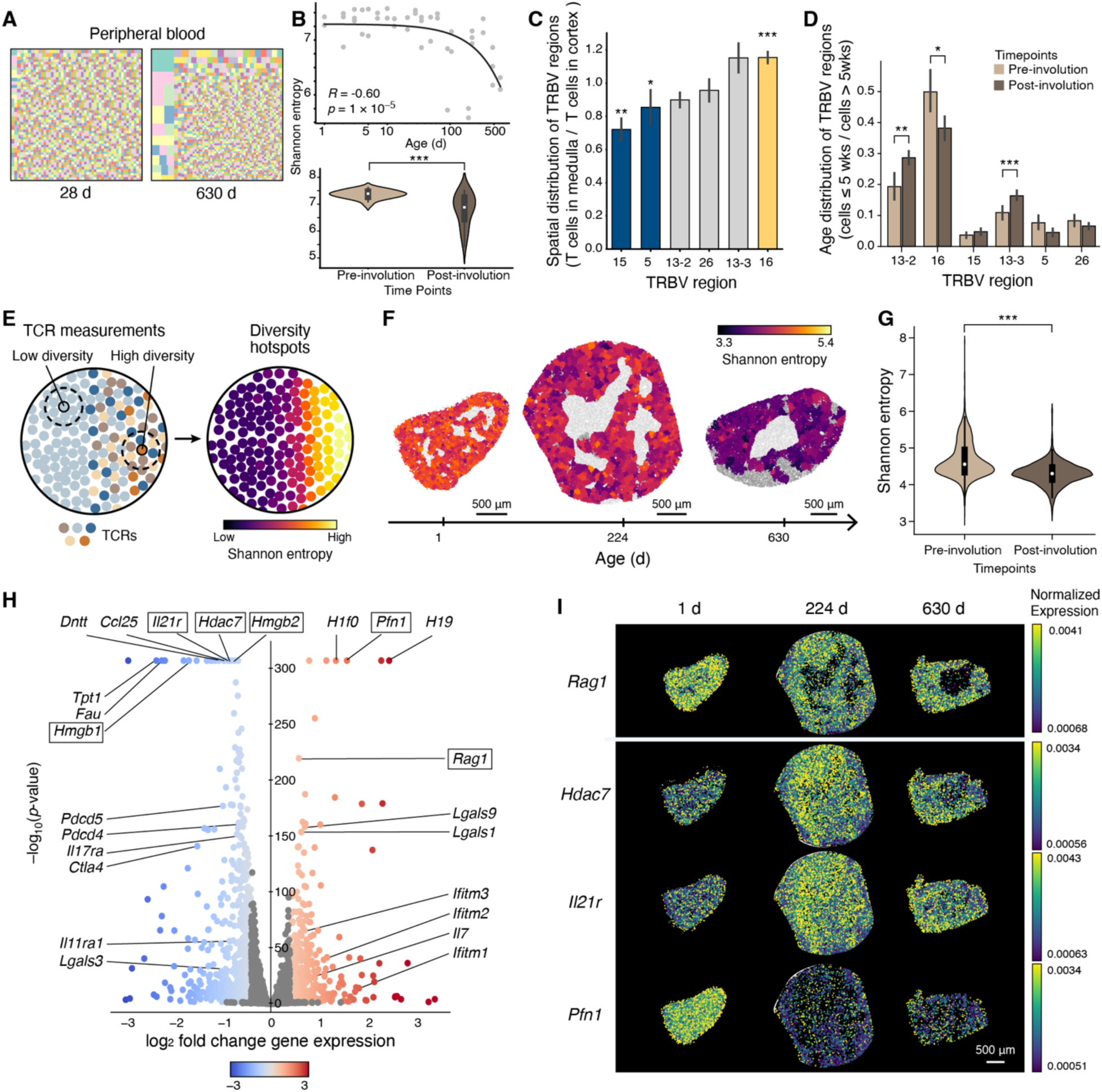
T cell receptor repertoire decline in thymus and periphery across age. **a,** TCR repertoire diversity in the peripheral blood using CDR3 amino acid sequences to define clonotypes, downsampled to 7500 cells per time point, at representative young (28 d) and aged (630 d) time points. Each colored square represents a unique clone, with the size of each square representing the fraction of the total repertoire belonging to each clone. **b,** TCR repertoire diversity in the spatial data using CDR3 regions, downsampled to 200 cells per time point, visualized individually (top), and separated by pre-involution (<= 35 d, n=12 mice) and post-involution (>35 d, n=9 mice) time points (bottom). **c,** Fraction of TRBV regions in medullary vs. cortical regions of the thymus across all time points (n=21 mice) in the spatial data. Two-sample paired t test and Benjamini-Hochberg correction used for comparison between TRBV regions. Error bars represent standard error. **d,** Fraction of TRBV regions in pre-involution (<= 35 d, n=12 mice) versus post-involution thymus (>35 d, n=9 mice). Two-sample t test and Benjamini-Hochberg correction used for comparison between age groups. Error bar represents 95% confidence interval. **e,** Schematic demonstrating spatial diversity measurement: for each T cell, the nearest 20 T cells are sampled, and then a Shannon entropy score is calculated. **f,** Tissue maps of T cell repertoire diversity, as quantified by Shannon diversity over three representative time points. **g,** Differences between Shannon diversity of T cell repertoires between pre-involution (<= 35 d, n=12 mice) and post-involution (>35 d, n=9 mice) thymus by two-sample t test . **h,** Differential gene expression between high (top 10%) and low diversity (bottom 10%) spatial regions of the thymus (n=21 mice). Color bar denotes log2 fold change. **i,** Visualization of selected differentially expressed genes across time points in aging thymus. Color bar denotes normalized gene expression. * = p < 0.05, ** = p < 0.005, *** = p < 0.0005.

To determine if spatial regions within the thymus exhibit biases in TCR recombination and selection, we analyzed the frequency of different T-cell receptor beta variable (TRBV) genes, which compose the variable (V) regions selected for during V(D)J recombination. Aggregating across all time points, we first found evidence that the TCR selection process is biased for certain V regions: *Trbv15* (p = 0.002, by two-sample t test) and *Trbv5* (p = 0.02, by two-sample t test) are biased for cortical regions involved in positive selection, while *Trbv16* (p = 2.6 x 10^-10^, by two-sample t test) is biased for medullary regions involved in negative selection. We did not observe statistically significant differences for J regions (**Fig. 4c and Extended Data Fig. 8a**). These results provide direct evidence of spatial variation in thymic selection pressure, which had previously only been inferred from single-cell studies^56^, reveal how dynamic changes in cortical-medullary structures over time may shape the TCR repertoire. Second, we identified biases with age by comparing pre- and post- thymic involution time points. Biases across TRBV and TRBJ included *Trbv13-2* (p = 0.005, by two-sample t test) and *Trbv13-3* (p = 0.003, by two-sample t test) enriched in aged, post-involution time points and *Trbv16* (p = 0.02, by two-sample t test) enriched in younger, pre-involution time points (**Fig. 4d and Extended Data Fig. 8b**). Last, to disentangle true age-related biases in V(D)J recombination from changes in the spatial structure of the thymus, we analyzed TCRs in the cortex and medulla separately and found these age-related shifts hold true (**Extended Data Fig. 8c**,d). Thus, biases in V(D)J recombination over aging cannot solely be attributed to differences in composition of cortex and medulla fraction or differences in CD4+ and CD8+ with age, as CD8+ T cells exhibit higher frequencies of both *Trbv16* and *Trbv13-357*.

We next examined how changes from thymic involution affect TCR diversity over time. We developed an analysis for quantifying diversity hotspots in our spatial data: for each bead containing a TCR CDR3 region, we sampled nearby TCRs to generate a spatial diversity score, the Shannon entropy of the sub-sampled TCRs (**Fig. 4e**). Interestingly, we found that not only does thymic size decrease with time, but the spatial diversity of the tissue worsens, as well; older thymi contain more “diversity dead zones” (p < 0.005 by two-sample t test, **Fig. 4f,g**). This finding raises the notion that therapies focusing on enlarging the thymus or increasing cellularity may not fully restore its function if compounding functional impairments, such as reduced TCR diversification, are not addressed.

### Differentially expressed genes in T cell diversification and differentiation

To investigate gene expression changes associated with these diversity hotspots and dead zones, we performed a niche-based differential expression analysis comparing gene expression differences between high-diversity and low-diversity regions. We identified several genes associated with high diversity, such as *Rag1*^58^, involved in V(D)J recombination, which was more highly expressed in hotspots (p < 0.001, Pearson’s r = 0.58) (**Fig. 4g**). In contrast, low-diversity regions exhibited expression of *Hmgb1* and *Hmgb2* (p < 0.005), high mobility group proteins shown to modulate chromatin architecture and regulate memory and exhausted T cell differentiation^59^. Their expression, along with the previously observed *Satb1+* increase with age, further implicate chromatin remodeling in age-related alterations of T cell developmental trajectories in the thymus.

Gene set enrichment analysis (GSEA) of genes in high-diversity regions revealed a gene set for epithelial-mesenchymal transition (EMT) (p = 1.54 x 10^-22^), involved in initiating thymic fibrosis and adipogenesis associated with aging and diminished regenerative capacity^60,61^ (**Extended Data Fig. 9a**). Conversely, low-diversity regions were enriched for an *Il-6/Jak/Stat3* signaling gene set (p = 7.2 x 10^-5^) (**Extended Data Fig. 9a**), indicating increased inflammation.

We found that *Dntt,* which encodes for terminal deoxynucleotidyl transferase (TdT), was highly expressed in low-diversity regions (p < 0.005). This was unexpected, as TdT is known for its role in adding N-nucleotides during V(D)J recombination, which typically increases the diversity of the CDR3 region^62–64^. To better understand this finding, we examined the expression of *Dntt* across individual cell types in low- vs. high-diversity niches and found that DP cells exhibited significantly higher *Dntt* expression in low-diversity regions (p < 0.005, **Extended Data Fig. 9b**). Further analysis revealed that *Dntt* expression increases with age in thymocytes, beginning in DN(Q) cells and through CD4+ and CD8+ stages (**Extended Data Fig. 9c**). We also observed a significant age-dependent increase in the *Dntt/Rag1* expression ratio in DP cells, despite decreased diversity (DP(P): p < 0.005, R = 0.42; DP(Q): p < 0.005, R = 0.30, **Extended Data Fig. 9d**). While TdT typically promotes diversity, its elevated expression in low-diversity regions could suggest an age-associated dysregulation in TdT activity. Existing literature shows that TdT enzymatic function decreases in aging thymuses in both humans and mice^65,66^. This diminished activity may impair the intended diversity-promoting role of TdT, leading to the formation of “diversity dead zones,” where despite high TdT expression, the actual diversity generated is limited. Another possibility is that TdT expression remains elevated in certain cell types, but the reduced recombination efficiency or altered selection pressure in aged thymuses could lead to the retention of cells with limited receptor diversity.

To better understand the gene expression in these low-diversity regions, we next aimed to identify genes contributing to diversity in an age-dependent manner. Visualized in the tissue context, genes like *Rag1* showed insignificant variation over age, in contrast with genes such as *Hdac7*, encoding an inhibitor of apoptosis^67^, *Pfn1*, encoding an actin-binding protein important in cell division^68^, and *Il21r*, encoding a receptor for IL-21, a cytokine improving thymic recovery in aging^69^, which exhibited dramatic shifts with age (**Fig. 4h**). Honing in on the T cell developmental trajectory, we evaluated the statistical significance of these shifts, confirming that while *Rag1* expression does not change with age, *Pfn1*, *H1f0*, an epigenetic regulator of cell proliferation^70^, *Il21r,* and *Hdac7* exhibited significant changes in developing thymocytes with age, such as increased *Il21r* in DP(P) cells with age (p = 2.43 x 10^-3^, Pearson’s r = 0.72), increased *Hdac7* in DP(P) cells with age (p = 4.79 x 10^-4^, Pearson’s r = 0.78) and decreased *H1f0* in DP(P) cells with age (p = 4.05 x 10^-4^, Pearson’s r = -0.79) (**Extended Data Fig. 9c**).

Beyond diversification, we were also able to use our approach to identify differentially expressed genes between CD4+ T and CD8+ T cell niches, identifying *Itm2a* (p < 0.005) as highly expressed in CD4+ T cell niches, which has been shown to be important for CD4+ SP commitment^71^, and *Runx3* (p = 0.0007) in CD8+ T cell niches, which is required for CD8+ fate commitment^72,73^ (**Extended Data Fig. 9e**). We found that while *Runx3* did not show changes in gene expression with age in developing T cells, *Itm2a* showed decreased expression in DP(P) (p = 2.51 x 10^-2^, Pearson’s r = -0.57) and CD4+T (p = 3.75 x 10^-2^, Pearson’s r = -0.55), highlighting an area for future study of changes in CD8+ SP commitment with age (**Extended Data Fig. 9f**). Together, we identified genes associated with T cell fate commitment and diversification through spatial analyses that could be leveraged for molecularly tailored thymus regeneration approaches.

## DISCUSSION

Thymic involution is a complex process that involves changes in the cellular composition and architecture of the thymic microenvironment. In this study, we used spatial methods to understand the critical cell interactions shaping this process, developing a framework for studying these changes over aging and their impact on immune function. To do so, we built a spatial and single-cell transcriptomic reference for thymic involution, identifying age-associated shifts in cellular organization and T cell repertoire diversification. Our analyses reveal substantial morphological shifts in the thymus over time, affecting cell composition, organization, and interactions. By coupling clonal T-cell identities *in situ* with spatial mapping of gene expression patterns, we linked molecular pathways involved in thymic involution to the decline of thymic output, as measured by T cell diversity. Using these data, we defined tissue organizational hallmarks of thymic involution in aging, which may serve as references for instances of thymic involution in other contexts, such as acute infection.

Furthermore, our analysis of the thymic microenvironment further revealed the complexity of cell types constituting the thymus and the breadth of interactions between stromal macrophages and other innate immune cells to support T cell differentiation. We describe an intercellular communication network between thymocytes and stromal macrophages closely tied to thymic involution. These findings collectively contribute to our understanding of immunological aging, hold promise for potential thymic regeneration strategies, and provide insights into the broader landscape of tissue changes over time.

Broadly, we have generated this resource and accompanying analyses as starting points for studying tissue dynamics over time scales of organism lifespan. While mouse immunity shares many aging features in common with humans, longitudinal human thymus samples will be needed to capture the diversity of human aging before interventions can be explored. Furthermore, our observation of the changes in gene expression of several chromatin-related and histone-related genes suggests the importance of including epigenomic readouts of tissue dynamics. Our findings will lastly serve as a resource for models of thymic regeneration to assess to what extent regeneration efforts recapitulate thymic function in healthy homeostasis.

## Data Availability

All data is available at the Broad Single Cell Portal, with accession SCP3197 (https://singlecell.broadinstitute.org/single_cell/study/SCP3197).

## Code Availability

Code to reproduce each figure is available at github.com/immunoliugy/

## MATERIALS AND METHODS

### Sample information and processing

Thymus samples for spatial Slide-TCR-seq were derived from C57BL/6J male mice (Jackson Laboratories). The thymus lobes were isolated, weighed, and embedded in optimal cutting temperature (OCT) compound. Following this, the specimens were promptly frozen on dry ice and stored at -80°C. The right lobes from 21 distinct time points were selectively chosen for spatial Slide-TCR-seq, as outlined in **Supplementary Table 1.**

Thymus and blood samples designated for single-cell RNA sequencing (scRNA-seq) were obtained from C57BL/6J mice (Janvier). These mice were housed under Specific and Opportunistic Pathogen-Free (SOPF) conditions, adhering to a 12-hour light: dark cycle, with ad libitum access to food and water. Animals aged between 4 and 90 weeks were carefully matched for age and co-housed within the same facility until organs were isolated. All animal procedures followed the institutional laboratory animal research guidelines and were approved by the governmental authorities (Regional Administrative Authority Karlsruhe, Germany).

### Histological processing

Serial sections, each 10 μm in thickness, were obtained from the frozen tissue samples and mounted onto glass plus microscope slides. For hematoxylin and eosin (H&E) staining, the Leica ST5010 Autostainer XL (Leica Biosystems) was used. Sections were immersed in xylene, sequentially processed through a graded ethanol series (100% and 95%), and then stained with hematoxylin. The nuclei were differentiated with mild acid treatment, followed by “bluing” through exposure to a weakly alkaline solution and subsequent water washing. Eosin staining was applied, and sections were again processed through a graded ethanol series (100% and 95%), xylene, dehydrated, and finally cover-slipped using the Leica CV5030 Fully Automated Glass Coverslipper. Brightfield images were captured with the Leica Aperio VERSA Brightfield, Fluorescence & FISH Digital Pathology Scanner, using the 20x objective.

### *In situ* transcriptome processing via Slide-seq

Slide-seq arrays were prepared, and spatial bead barcodes were sequenced following Slide-seqV2 74 protocol. Custom-synthesized barcoded beads featured a sequence (5’- TTT_PC_GCCGGTAATACGACTCACTATAGGGCTACACGACGCTCTT CCGATCTJJJJJJJJTCTTCAGCGTTCCCGAGAJJJJJJNNNNNNNVVT30-3’) with a photocleavable linker (PC), a bead barcode sequence (J, 14 bp), a UMI sequence (NNNNNNNVV, 9 bp), and a poly dT tail.

OCT-embedded frozen tissue samples were warmed to -20°C in a cryostat (Leica CM3050S) and serially sectioned at a 10 μm thickness (2-3 Slide-seq array replicates per sample), with consecutive sections used for hematoxylin and eosin staining and immunofluorescence staining. Each tissue section was affixed to an array and moved into a 1.5 mL Eppendorf tube for downstream processing. The sample library was prepared as previously described ^74^, including a library amplification via a PCR program of 1 cycle of 98°C for 2 min, 4 cycles of 98°C for 20 s, 65°C for 45 s, 72°C for 3 min, 9 cycles of 98°C for 20 s, 67°C for 20 s, 72°C for 3 min, and 1cycle of 72°C for 5 min. The PCR was performed in a final volume of 200 mL of PCR mix, divided into 4 PCR tubes. Libraries were sequenced using the following read structure on a NovaSeq (S4 or X; Illumina): Read1: 42 bp; Read2: 41 bp; Index1: 8 bp, and sequences were processed as previously described ^74^ using the pipeline available at (https://github.com/MacoskoLab/slideseq-tools).

### TCR clonotype enrichment for spatial TCR sequencing

For mouse TCRs, 56 PCR primers specific to the V segments of mouse alpha and beta TCR genes **(Supplementary Table 2)** were used. The alpha and beta primers were separately pooled at a final concentration of 100 μM, combining 5 μL of each primer at 100 μM concentration. The PCR reaction was prepared as follows: 10 ng of Slide-seq cDNA library, 0.25 μL of 100 μM T7-TCRV primer pool, 12.5 μL of KAPA Hifi Hotstart Readymix 2X, and Ultrapure water up to 25 μL. The following program was used on the thermal cycler: 5 cycles of 95°C for 5 min, 65°C for human primer pool/70°C for mouse primer pool for 30 sec, and 72°C for 3 min, followed by a hold at 4°C. Subsequently, 25 μL of Ultrapure water was added with 30 μL of SPRIselect or AMPure XP beads for 0.6X clean-up, following manufacturer’s instructions, and eluted into 9 μL of water.

The elution was used in a subsequent PCR prepared as follows: 9 μL of eluted product, 0.5 μL of 100 μM Truseq-P5 Hybrid primer (5’ AATGATACGGCGACCACCGAGATCTACACTCTTTCCCTACACGACGCTCTTCCGATC T 3’), 0.5 μL of 100 μM T7 PCR primer (5’ TCTAGATAATACGACTCACTATAGGG 3’), 12.5 μL of KAPA Hifi Hotstart Readymix 2X, and Ultrapure water up to 25 μL. The following program was used on the thermal cycler: 1 cycle of 98°C for 2 min, 10 cycles of 98°C for 1 min, 60°C for 30 sec, and 72°C for 3 min, 1 cycle of 72°C for 5 min, and hold at 4°C. Upon completion, 25 μL of Ultrapure water was added with 30 μL of SPRIselect or AMPure XP beads for 0.6X clean-up, following the manufacturer’s instructions, and eluted into 8 μL of water.

The product was primarily amplified by an in vitro transcription reaction (IVT). The reaction included 8 μL of the sample, 10 μL of NTP Buffer Mix, and 2 μL T7 RNA Polymerase Mix, following the HiScribe RNA Synthesis kit instructions. The reaction was incubated at 37°C for 2 hours. Upon completion, 30 μL of Ultrapure water was added with 30 μL of SPRIselect or AMPure XP beads for 0.6X clean-up, as per the manufacturer’s instructions, and eluted into 20 μL of water.

Subsequently, 20 μL of the product was transferred to a reverse transcription (RT) reaction. This included 40 μL of Maxima 5X RT buffer, 20 μL of 10 mM dNTPs, 5 μL of RNAse inhibitor, 2 μL of 100 μM Truseq-P5 Hybrid primer, 10 μL of Maxima H-RTase, and Ultrapure water up to 200 μL. The reaction was incubated at 42°C for 2 hours. Then, 120 μL of SPRIselect or AMPure XP beads were added for a 0.6X clean-up following the manufacturer’s instructions, and the elution was performed into 20 μL of water.

From this, 100 ng of DNA was taken for a subsequent index PCR. This was added to the following reaction mix: 100 μL of KAPA Hifi Hotstart Readymix 2X, 4 μL of 100 μM Truseq-P5 Hybrid PCR primer, 4 μL of 100 μM Nextera PCR primer (i7), and Ultrapure water up to 200 μL. The reaction was divided into 4 PCR tubes, each containing 50 μL (25%) of the total volume. The following program was used on the thermal cycler: 1 cycle of 98°C for 2 min, 10 cycles of 98°C for 1 min, 67°C for 20 sec, and 72°C for 3 min, 1 cycle of 72°C for 5 min, and a hold at 4°C. The four parts were recombined, and 120 μL of SPRIselect or AMPure XP beads was added for a 0.6X clean-up following the manufacturer’s instructions, eluting into 10 μL of water. 1 µL of each library was used to confirm the fragment size distribution and yield using an Agilent High Sensitivity DNA Kit (Agilent Technologies) on an Agilent 2100 bioanalyzer.

### Dissociated single-cell RNA/TCR sequencing and data preprocessing

Heart blood was obtained through cardiac puncture in mice following cervical dislocation and stored in 0.5% EDTA for subsequent processing. Thymus lobes from mice sacrificed by cervical dislocation were isolated, washed in PBS, and passed through a 70 μm cell strainer. Red blood cells were lysed using ACK lysis buffer (Thermo). Freshly isolated immune cells post-red blood cell lysis were blocked with rat anti-mouse CD16/32 (0.5 μg per well, eBioscience). Subsequently, respective antibodies in PBS were added in a total volume of 50 μL and stained for 30 minutes. The antibodies used included CD45-BV510, CD3-FITC, CD11b-PE-Dazzle. eFluor 780 fixable viability dye (eBioscience) was employed per the manufacturer’s protocol to exclude dead cells.

For scRNA/TCR-Seq, cells were divided into 8 aliquots per animal sample and pre-incubated for 10 minutes with titrated amounts of TotalSeq™ hashtag antibodies, C0301-C0308 (BioLegend). Cells were sorted on a BD Aria Fusion cell sorter using a 100 μM nozzle and 4-way purity mode. From peripheral blood, viable T cells (live, CD45+, CD3+) were sorted in 20 μL 0.04% BSA in PBS and kept on ice until processing. From thymus samples, all viable cells (live) were sorted in 20 μL 0.04% BSA in PBS and kept on ice until processing.

Single-cell capture, reverse transcription, and library preparation were conducted on the Chromium platform (10x Genomics) with the single-cell 5ʹ reagent v2 kit (10x Genomics) following the manufacturer’s protocol, using 40,000 cells as input per channel. Each pool of cells underwent library quality testing, and library concentration was assessed. The final library for each pool was subjected to paired-end sequencing (26 and 92 bp) on one Illumina NovaSeq 6000 S2 lane. Raw sequencing data were processed and aligned to the mouse genome (GRCm39 - mm39) using the CellRanger pipeline (10x Genomics, version 7.1.0).

### TCR clonotype enrichment and processing of dissociated TCR sequencing

Following the preparation of single-cell suspensions and T cell sorting as described above, 40,000 antibody-hashed cells per animal were loaded onto a 10x Chromium controller (Chip G) per the manufacturer’s protocol using the Chromium Single Cell V(D)J Reagent Kits (v1.1 Chemistry) and featured barcoding technology for cell surface proteins to capture antibody hashtags. The resulting emulsion was recovered and immediately subjected to reverse transcription. Subsequent steps included cDNA isolation and library preparation, adhering to the manufacturer’s protocol. Each of the final V(D)J libraries underwent paired-end sequencing (26 and 92 bp) on a single Illumina NovaSeq 6000 S2 lane. Raw sequencing data underwent processing and alignment to the mouse genome (GRCm39 - mm39) using the CellRanger pipeline (10x Genomics, version 7.1.0).

Seurat datasets were generated for blood and thymus at each time point. Singlets were identified per the published Seurat vignette (https://satijalab.org/seurat/articles/hashing_vignette.html) and used for downstream analyses. Additionally, only cells with more than 500 and less than 4000 unique features were detected, and less than 5% of mitochondrial counts were considered for further analysis. Using scRepertoire AB-VDJ, information was added per sample using default settings, employing the CDR3 amino acid (”aa”) sequence for clonotype calling. The final datasets post-QC yielded 104,896 cells (peripheral blood) and 73,558 cells (thymocytes), respectively.

### Integration, clustering, and cell type identification of dissociated single-cell TCR sequencing data

Blood and thymus datasets underwent merging and integration using the harmony package, following the guidelines in the published vignette (https://portals.broadinstitute.org/harmony/SeuratV3.html). The integration process involved the application of the following arguments: NormalizeData(), FindVariableFeatures(selection.method = “vst”, nfeatures = 2000), ScaleData(), RunPCA(), RunHarmony(”orig.ident”, plot_convergence = TRUE, dims.use = 1:20), RunUMAP(reduction = “harmony”, dims = 1:20), FindNeighbors(reduction = “harmony”, dims = 1:20), FindClusters(resolution = 0.5). After removing cells failing quality control, integration was reiterated with the same settings, and final transcriptional clusters for downstream analyses were identified using the FindClusters function (resolution = 0.7).

The dissociated thymocyte dataset underwent reference annotation using the SingleR package (…,de.method=”wilcox”) and a published reference dataset^33^ for each individual sampling time point. Annotation of mouse peripheral blood datasets was carried out manually, based on the final transcriptional clustering post quality control and integration, using canonical marker genes (**Extended Data Fig. 1b**) and differential gene expression analyses through the FindAllMarkers function in R.

### Differential abundance testing

The statistical significance of changes in cell type abundance over time in the single-cell data was assessed using Pearson correlation through the cor.test(…, method= “pearson”) function. For visualization, log2 fold changes of cell type proportions at each time point relative to the 4-week time point were computed and graphically represented. Cell types exhibiting a significant correlation with increasing age (p < 0.05) were color-annotated. Density visualization in scanpy ^75^ was performed following the conversion of the final Seurat objects to anndata objects using the SeuratDisk function (Convert(…,, dest = “h5ad”)). Cell density plots were generated in Python using the scanpy ^75^ function sc.tl.embedding_density(…, basis=’umap’, groupby=’week’).

### VDJ analysis of peripheral blood

For VDJ analysis, only cells with productive TCRs were considered, and the datasets were downsampled to 2500 cells (Thymus) / 7500 cells (Blood) per time point using scRepertoire. The VDJ information for the downsampled objects was updated with the same settings as previously described. To visualize the occupied repertoire per cluster at each time point, the occupiedscRepertoire() function was employed. Diversity calculation was performed using clonalDiversity(…, cloneCall = “aa”, group.by = “orig.ident”, n.boots = 100, exportTable = T, return.boots = F). Clonal homeostasis was depicted using clonalHomeostasis(…, cloneCall = “aa”). Treemaps were generated for the top 2500 TCR clonotypes per sample and time point, based on the cellranger output (clonotypes.csv), using the treemap(…, type=“index”) function.

### Unsupervised clustering for region identification of cortex and medulla

To delineate cortex and medulla regions in the thymus, we initiated the process by manually selecting the entire tissue, and removing points outside the primary tissue region using the matplotlib ^76^ Lasso Selector tool. Subsequently, standard data pre-processing steps were implemented, which involved filtering out beads with fewer than 50 genes, excluding genes present on fewer than 3 cells, eliminating beads with over 5% mitochondrial reads, normalizing, and log-transforming the data. Unsupervised clustering was then performed using scanpy ^75^.

Clusters were annotated based on the top twenty differentially expressed genes if they contained cortex markers (*Themis, Top2a, Npm1, Rag1, Dntt*) or medulla markers (*Tmsb4x, Krt5*). Unassigned points were then labeled as either cortex or medulla depending on the majority of their nearest neighbors within a 30 μm distance. This process was reiterated to capture the remaining unassigned cells at a 50 μm distance.

### Thymic medullary surface area to perimeter ratio calculation

The surface area-to-perimeter ratio of thymic medullary regions was computed through the following steps. Initially, the surface area of the medulla was determined by quantifying the total number of beads in regions previously annotated as the medulla. Subsequently, the perimeter of the medulla was defined as beads that, within a 50 μm radius, had more than half of their neighboring beads annotated as cortex. The division of these two values across different time points facilitated the calculation of the medullary surface area-to-perimeter ratio.

### Applying RCTD for cell type assignment on spatial data and calculating cell type proportions

Cell type assignment for our spatial thymus data was accomplished using Robust Cell Type Decomposition (RCTD)^34^ in doublet mode, employing a single-cell mouse thymus reference^23,33^. The cell type proportions in **Fig. 2a** were computed using the top cell weight derived from RCTD. For the grouped cell type labels, Effector Lymphocytes = CD4+T, CD8+T, Treg, IELpA, IELpB_NKT, γδT, αβT(entry), NK; Thymocytes = DN(P), DN(Q), DP(P), DP(Q); Stroma = VSMC, Ery, Epi_unknown, Fb, TEC_early, Endo, HSC; APC = B, DC1, DC2, pDC, Mac, Mono; and TEC =either cTEC or mTEC, depending on the plotted region.

### Comparing T cell and thymocyte detection in single-cell and spatial data

Cell types classified as T cell/thymocyte types included CD4+ T, CD8+ T, DN(Q), DP(P), DN(P), αβT(entry), γδT, and Treg. Following this, the proportion of cells corresponding to these cell types was computed for the scRNA-seq data. In the spatial data analysis, the top cell weight from RCTD was used to label cell types, which were then used to identify the fraction of cells matching T cell/thymocyte types.

### TCR identification in spatial data and spatial diversity analysis

Variable TCR sequences were identified using MiXCR (v3.0.13) ^77^ to align to the mouse genome. For MiSeq data, accounting for sequencing errors-introduced diversity involved correcting the CDR3 amino acid sequences with a Hamming distance of 1 and collapsing to the most predominant sequence. Subsequently, we identified the bead barcode and UMI sequence, eliminating all reads without a match within a Hamming distance of 1 to an in situ sequenced bead barcode.

In spatial diversity analyses, for each TCR, the nearest 20 neighboring TCRs were sampled. Shannon entropy was then used to calculate a diversity score for each bead. To address gaps in the capture, imputation was carried out through two rounds of dilation, expanding the beads in a 20 μm radius to fill the cortical space. Following imputation, differential gene expression analyses were conducted on the top 10th percentile and the bottom 10th percentile of beads based on diversity. Gene set enrichment analysis was performed using the MSigDB Hallmark 2020 collection.

### Quantifying spatial organization via interaction count statistic

For each pair of cell types at each time point, a z-score was computed by comparing the observed frequency of cell interactions for the pair within a 50 μm radius to the null hypothesis of the frequency of cell interactions for the same pair within a 150 μm radius. Subsequently, these z-scores were plotted over time, and Pearson’s r was employed to assess the strength of the correlation between changing z-scores and age.

### Receptor-ligand analysis

We used the Squidpy ^43^ integration of CellPhoneDB ^44^ and Omnipath ^45^ to identify shifts in receptor-ligand interactions at each time point. Subsequently, the mean values of each receptor-ligand cell-cell pair across time were employed to calculate Spearman’s R correlation and a p-value. After correcting the p-values through Bonferroni correction, unique interactions were categorized into those decreasing and increasing with age for visualization.

### Predicting age from spatial transcriptomic data

We first tokenized the raw single cell digital gene expression data using Geneformer ^78^ to identify genes that are more important to focus on to optimize predictive accuracy of the linear regression model. In order to apply Geneformer to mouse single-cell datasets, we extracted mouse gene to human ortholog gene mapping information from Ensembl BioMart ^79^ (version: Ensembl Genes 111, dataset: Mouse genes (GRCm39)) and mapped the mouse gene ensembl ids to their human orthologs. We then tokenized the mouse dataset and fed it into a 6-layer pretrained Geneformer model and extracted cell embedding from the second to last layer of the model to use as features for the linear regression model. In addition to the extracted cell embedding, a cell neighbors vector was concatenated for each cell, which sampled the nearest 100 cells and reflected the fraction of neighboring cell types. The linear model was trained on all mouse aging time points, and predictions were made for perturbations.

## Supplementary Materials

Supplementary Tables 1-2

Extended Data Figs. 1-9

## Acknowledgements

We thank Sandeep Kambhampati for helpful conversations. We thank Leslie Gaffney for figure editing. We thank Michael Kilian for supporting single-cell sequencing experiments. We thank the Broad Institute Flow Core for histology assistance. This study was supported by Impetus Grants, BroadIgnite, and the Deutsche Forschungsgemeinschaft (DFG, German Research Foundation; reference FR 4701/1-1). S.L. is supported by the NIH Molecular Biophysics Training Grant (NIH/NIGMS T32GM008313) and the National Science Foundation Graduate Research Fellowship. This work was supported by the National Institutes of Health (grant nos. R01HG010647 to F.C.). F.C. also acknowledges support from the Searle Scholars Award, the Burroughs Wellcome Fund CASI award, and the Merkin Institute. This work was supported by the Milky Way Research Foundation (to F.Z.).

## Author contributions

S.L., M.J.F., and R.R. designed the experiments; S.L, R.R., L.A.M. performed Slide-TCR-seq experiments; S.L., D.G., B.C., Q.G., and N.K. analyzed the data. M.J.F. performed single-cell and in vivo experiments; K.G., D.S., gave input on the manuscript and analyses; F.C. and F.Z. supervised the project with support from R.K.M.; S.L. wrote the manuscript with contributions from all authors.

