## Supplementary Figures for "Tissue architecture dynamics underlying immune development and decline in the thymus"

**Supplementary Materials**

Supplementary Tables 1-2
Extended Data Figs. 1-9

### Extended Data Figures

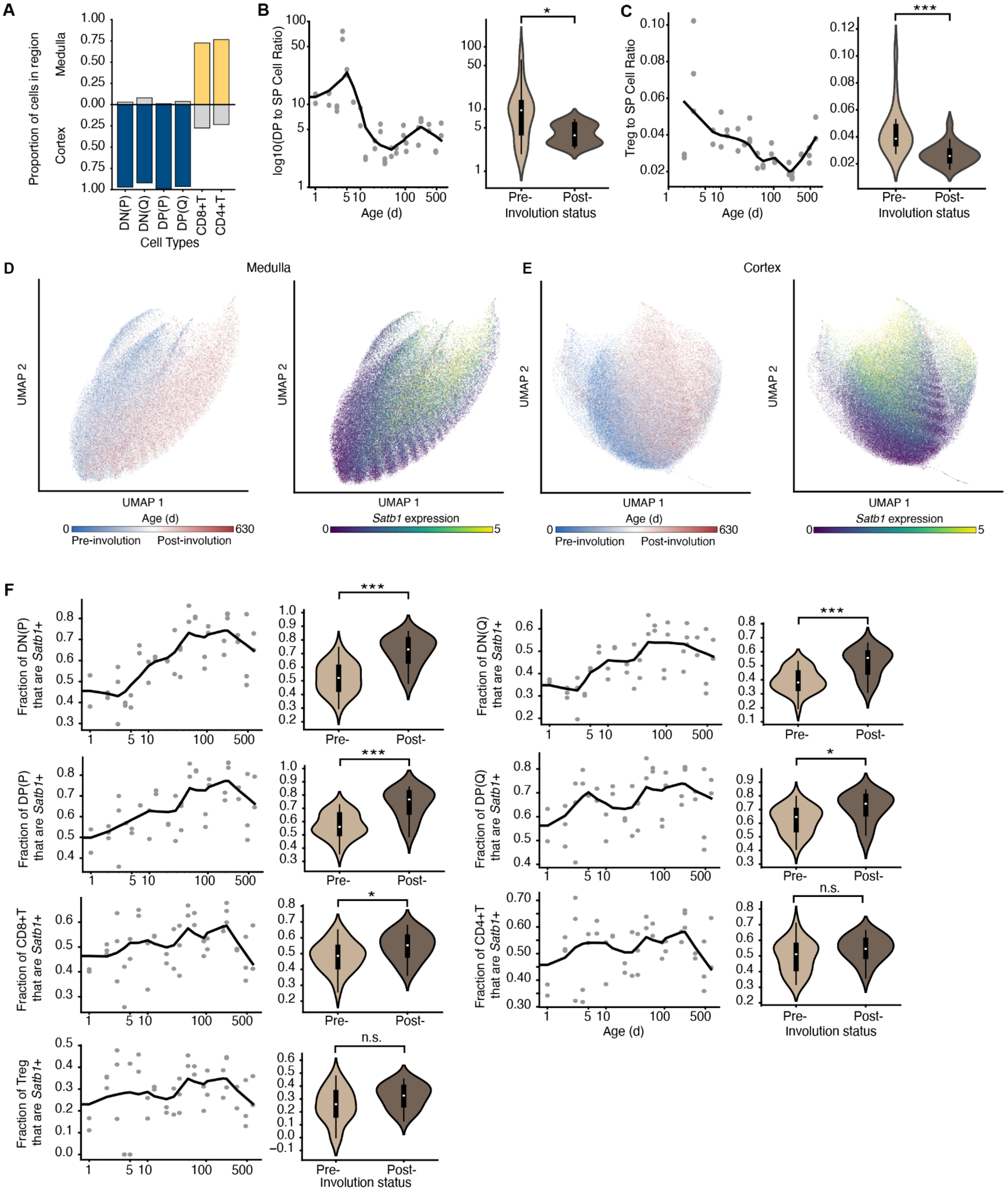


### Extended Data Fig. 1. Thymocyte population shifts with age

**a,** Proportion of developing thymocytes and T cells in cortical and medullary regions.

**b,** Ratio of DP to SP thymocytes visualized over all time points with local regression curve (left) and separated by pre-involution (<=5 wk, n=12 mice) and post-involution (>5 wk, n=9 mice) time points (right). Two-sample t test used to compare pre- and post-involution.
**c,** Ratio of Treg to SP thymocytes visualized over all time points with local regression curve (left) and separated by pre-involution (<=5 wk, n=12 mice) and post-involution (>5 wk, n=9 mice) time points (right). Two-sample t test used to compare pre- and post-involution.
**d,** Uniform Manifold Approximation and Projection (UMAP) of cells in medulla from spatial data colored by age (left) and *Satb1* expression (right).
**e,** Uniform Manifold Approximation and Projection (UMAP) of cells in cortex from spatial data colored by age (left) and Satb1 expression (right).
**f,** Fraction of developing thymocytes and T cells that are *Satb1*+ over all time points plotted individually and as a grouped comparison between pre- and post-involution time points. Two-sample t test used to compare pre- and post-involution, with Benjamini-Hochberg correction for multiple-hypothesis testing.

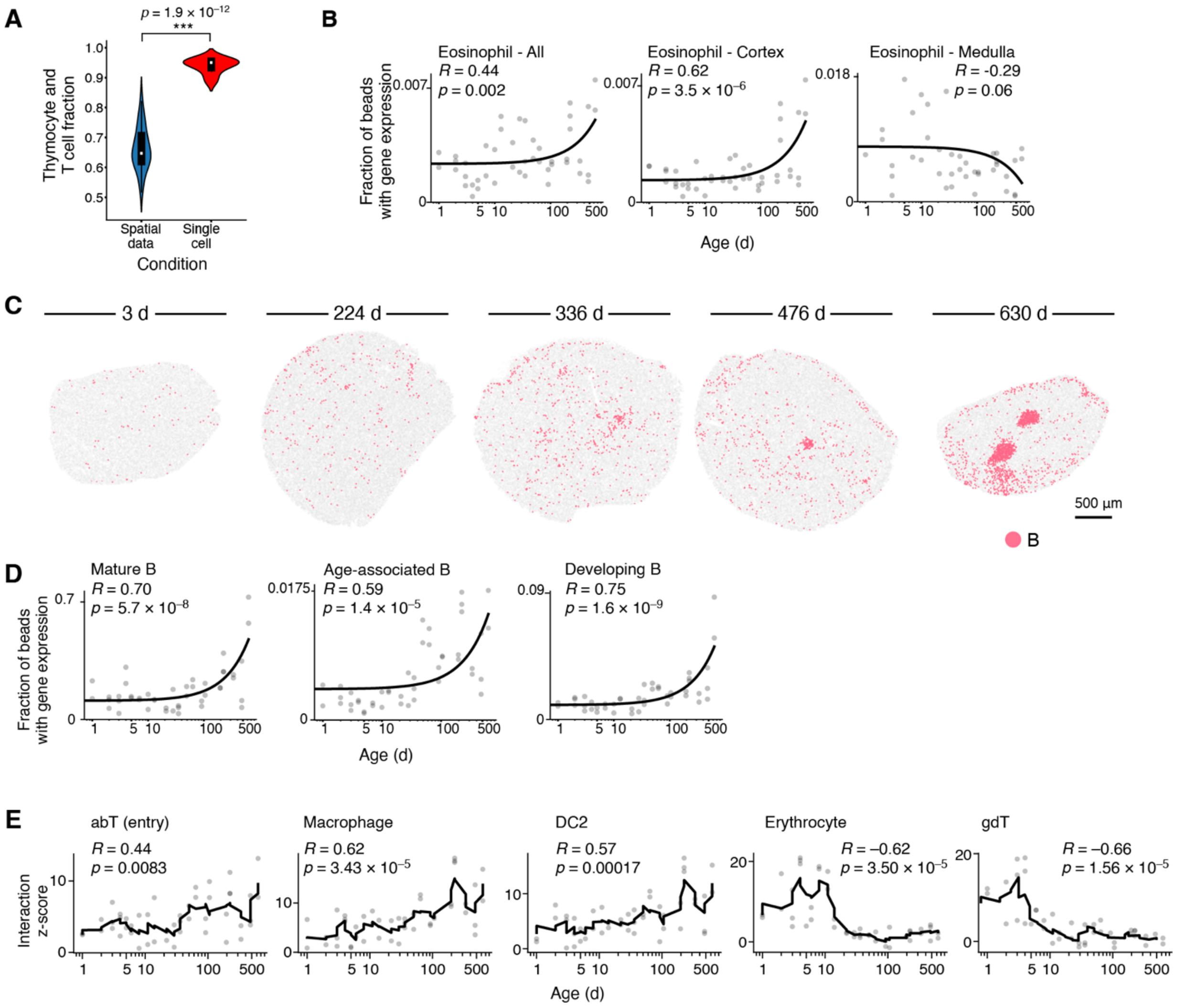


**Extended Data Fig. 2. Changes to the thymic cell populations and clustering in spatial data.**

**a,** Fraction of cells that are thymocytes or T cells in spatial (n=47) vs. single-cell (n=7) data across all samples. Two sample t-test used for comparison. Error bar represents 95% confidence interval.

**b,** Proportion of cells expressing eosinophil markers (*Siglecf, Epx, Clc, Rnase2, Ecp, Prg2*) with age in spatial data, separated by spatial location. Line of best fit plotted and Pearson's r used to calculate correlation over time.

**c,** Representative spatial maps of B cell aggregates in the thymus with age. Colored beads denote B cells assigned via RCTD.

**d,** Proportion of cells expressing mature B cell (*Igkc, Ms4a1, Cd20, Cd19, Cd22, Cd27*), age-associated B cells (*Itgax, Tbx21*), or developing B cell (*Vpreb1, Vpreb3, Ebf1, Pax5*) markers with age in spatial data, separated by spatial location. Line of best fit plotted and Pearson's r used to calculate correlation over time.

**e,** Significant changes in spatial self-clustering of cell-types over time measured via permutation testing, with a magnitude of $R > 0.5$ and p values corrected using Benjamini-Hochberg correction.

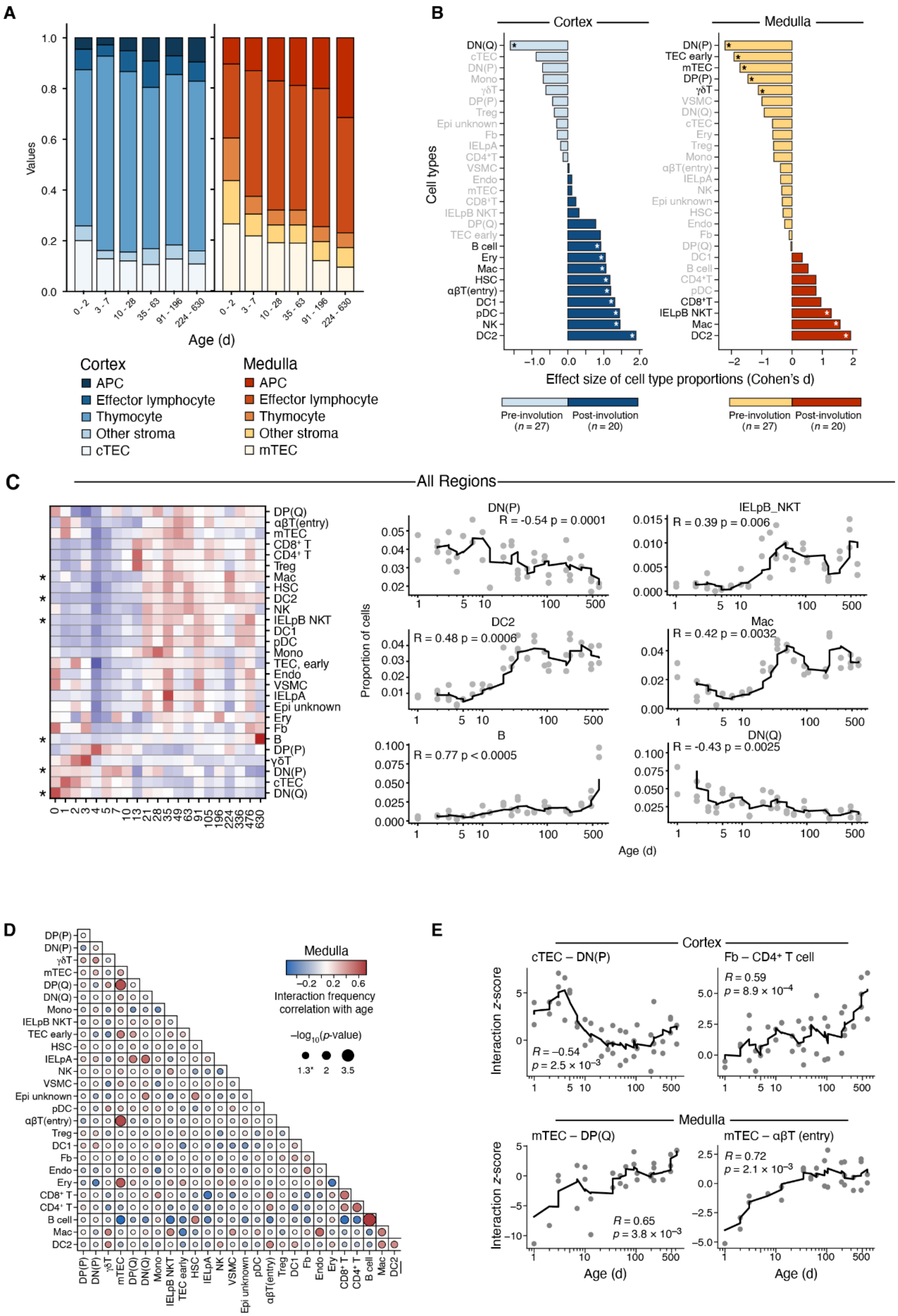


**Extended Data Fig. 3. Cell-cell interaction dynamics with age**

**a,** Cell type proportions over time in the cortex and medulla of antigen-presenting cells (APCs), effector lymphocytes, thymocytes, cortical and medullary thymic epithelial cells (cTEC, mTEC), and other stroma. Time points are grouped from **Fig. 2a** to have multiple mice per time range (0-2: n=7 spatial arrays over 3 mice; 3-7: n= 9 spatial arrays over 4 mice for cortex, 7 spatial arrays over 3 mice for medulla; 10-28: n=6 spatial arrays over 3 mice; 35-63: n=7 spatial arrays over 3 mice; 91-196: n=6 spatial arrays over 3 mice; 224-630: n=10 spatial arrays over 4 mice).
**b,** Changes in cell type composition in the spatial thymus data from Slide-TCR-seq in the cortex and medulla. Pre-involution time points are <= 5 weeks, while post-involution includes time points >5 weeks. * = $p < 0.05$.
**c,** Cell-cell interaction frequency correlation (Pearson's r) with age in all regions of the thymus. Positive correlation values indicate increase in interaction with age. Circle size denotes fdr adjusted p-value.
**d,** Changes to selected cell-type interactions in the cortex and medulla with age. Rolling average line plotted, window = 5. P-value denotes significance of Pearson's correlation with age.

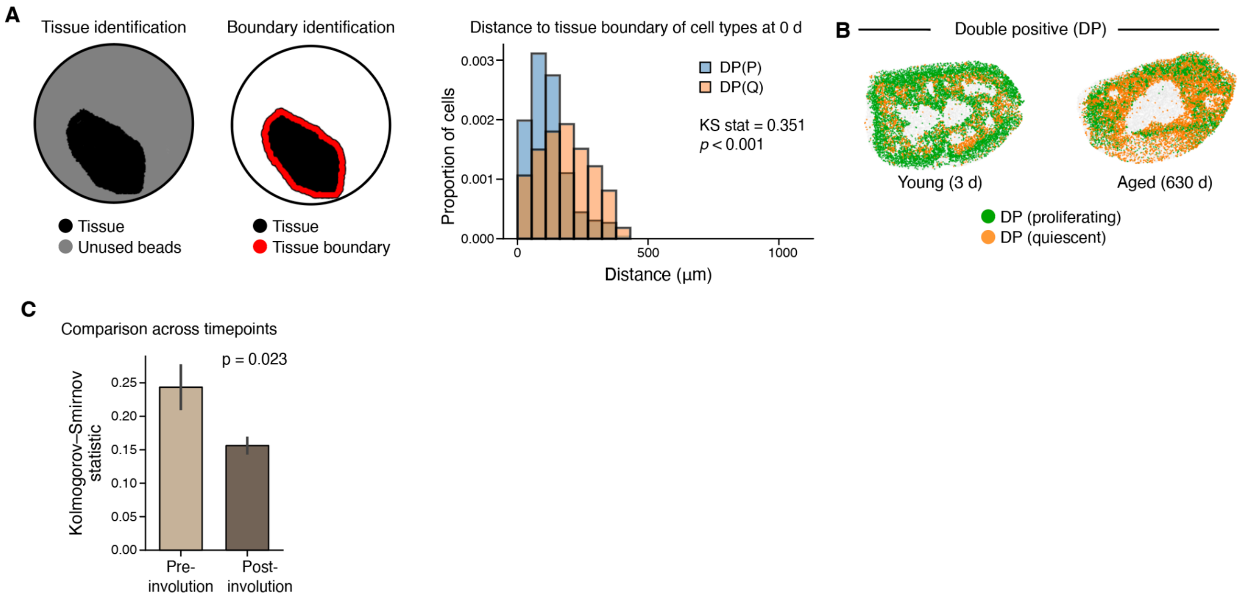


**Extended Data Fig. 4. Age-associated changes in spatial niches of DP(P) and DP(Q) cells**

**a,** Schematic demonstrating the analysis: first, tissue regions are identified. Second, the boundary is identified. Last, a distribution for the distance to the boundary is determined for each cell type and the Kolmogorov-Smirnov statistic is determined for comparing the distributions.
**b,** Slide-TCR-seq spatial arrays of representative young (3 d) and aged (90 wk) mouse thymi showing spatial organization of T cell precursors, double positive proliferating, (DP(P)) and double positive quiescent (DP(Q)) cell types.
**c,** Comparison of changes to DP(P) and DP(Q) organization with age by permutation test. Error bar represent standard error.

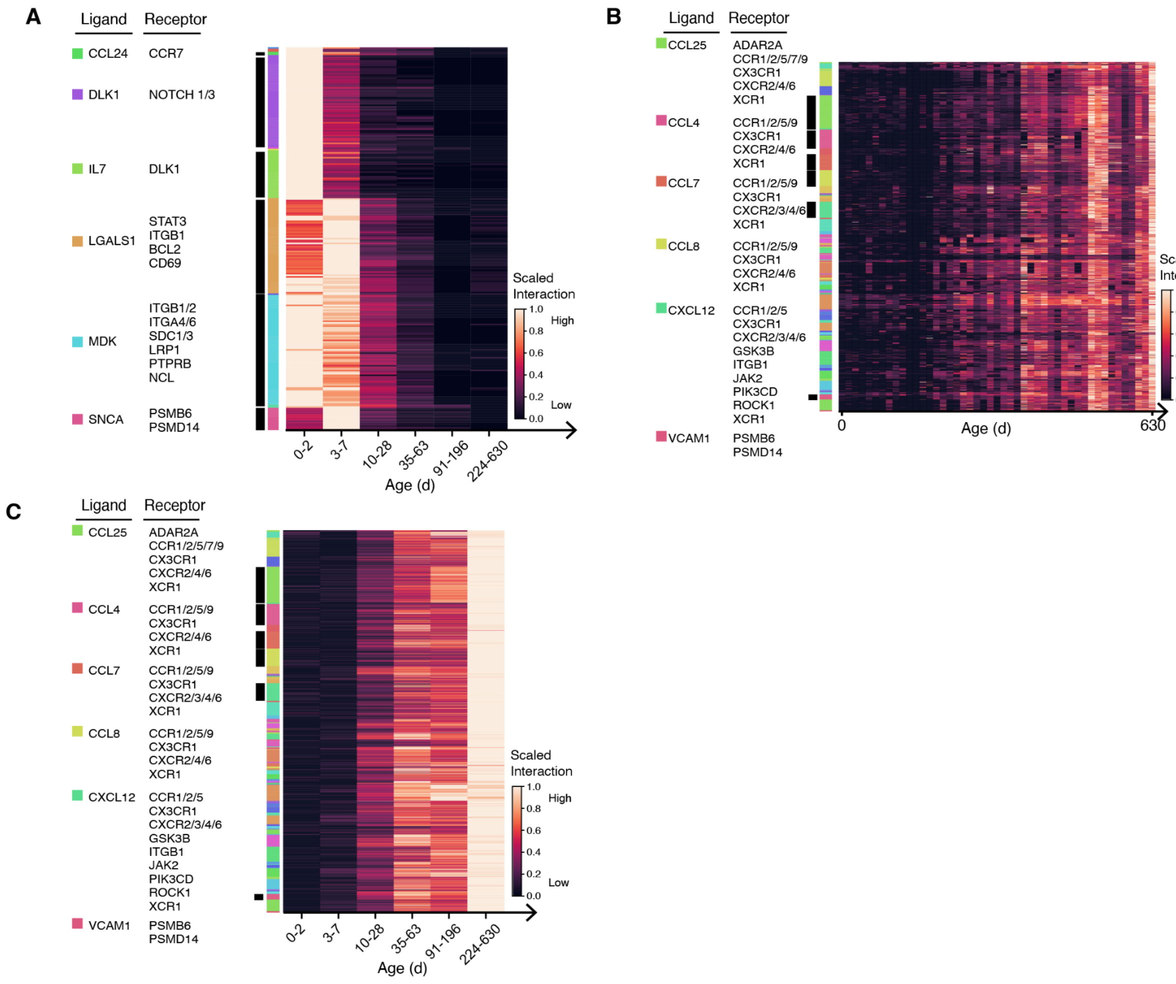


**Extended Data Fig. 5. Receptor-ligand interaction dynamics with age**

**a,** Decreasing receptor-ligand interactions with age; data from **Fig. 3b** is grouped into six age groups (0-2: n=7 spatial spatial arrays over 3 mice; 3-7: n=9 spatial arrays over 4 mice; 10-28: n=6 spatial arrays over 3 mice; 35-63: n=7 spatial arrays over 3 mice; 91-196: n=6 spatial arrays over 3 mice; 224-630: n=10 spatial arrays over 4 mice). Each row is a unique cell type-cell type receptor-ligand pair and is annotated by common receptor-ligands.

**b,** Increasing receptor-ligand interactions with age. Each row is a unique cell type-cell type receptor-ligand pair and is annotated by common receptor-ligands.

**c,** Increasing receptor-ligand interactions with age, data from (**b**) is grouped into six age groups (0-2: n=7 spatial arrays over 3 mice; 3-7: n=9 spatial arrays over 4 mice; 10-28: n=6 spatial arrays over 3 mice; 35-63: n=7 spatial arrays over 3 mice; 91-196: n=6 spatial arrays over 3 mice; 224-630: n=10 spatial arrays over 4 mice). Each row is a unique cell type-cell type receptor-ligand pair and is annotated by common receptor-ligands.

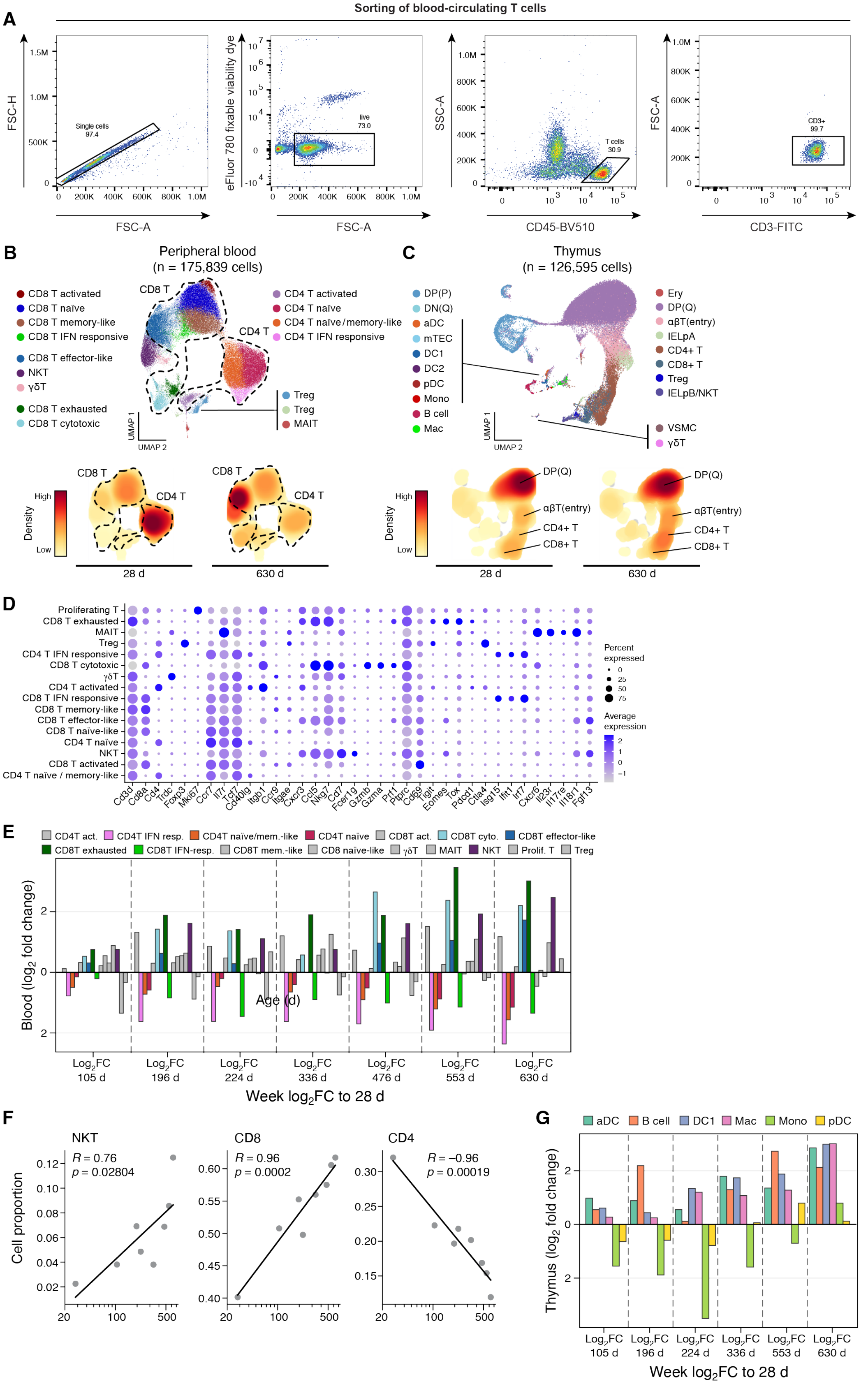

A
Sorting of blood-circulating T cells
Single cells 97.4
live 73.0
T cells 30.9
CD3+ 99.7
FSC-H
FSC-A
eFluor 780 fixable viability dye
SSC-A
CD45-BV510
CD3-FITC
B
Peripheral blood
(n = 175,839 cells)
CD8 T activated
CD8 T naïve
CD8 T memory-like
CD8 T IFN responsive
CD8 T effector-like
NKT
γδT
CD8 T exhausted
CD8 T cytotoxic
CD4 T activated
CD4 T naïve
CD4 T naïve/memory-like
CD4 T IFN responsive
Treg
Treg
MAIT
CD8 T
CD4 T
UMAP 1
UMAP 2
Density
High
Low
28 d
630 d
C
Thymus
(n = 126,595 cells)
DP(P)
DN(Q)
aDC
mTEC
DC1
DC2
pDC
Mono
B cell
Mac
Ery
DP(Q)
αβT(entry)
IELpA
CD4+ T
CD8+ T
Treg
IELpB/NKT
VSMC
γδT
D
Proliferating T
CD8 T exhausted
MAIT
Treg
CD4 T IFN responsive
CD8 T cytotoxic
γδT
CD4 T activated
CD8 T IFN responsive
CD8 T memory-like
CD8 T effector-like
CD8 T naïve-like
CD4 T naïve
NKT
CD8 T activated
CD4 T naïve / memory-like
Percent expressed
Average expression
E
CD4T act.
CD4T IFN resp.
CD4T naïve/mem.-like
CD4T naïve
CD8T act.
CD8T cyto.
CD8T effector-like
CD8T exhausted
CD8T IFN-resp.
CD8T mem.-like
CD8 naïve-like
γδT
MAIT
NKT
Prolif. T
Treg
Blood (log2 fold change)
Age (d)
Log2FC 105 d
Log2FC 196 d
Log2FC 224 d
Log2FC 336 d
Log2FC 476 d
Log2FC 553 d
Log2FC 630 d
Week log2FC to 28 d
F
NKT
R = 0.76
p = 0.02804
CD8
R = 0.96
p = 0.0002
CD4
R = –0.96
p = 0.00019
Cell proportion
G
aDC
B cell
DC1
Mac
Mono
pDC
Thymus (log2 fold change)
Week log2FC to 28 d

**Extended Data Fig. 6. Age-associated cell type changes in single-cell RNA-seq of peripheral blood and thymus**

**a,** Representative gating strategy for fluorescence-activated cell sorting (FACS) of blood-circulating T cells for dissociated single-cell RNA/TCR-seq.
**b-c,** (Top) Uniform Manifold Approximation and Projection (UMAP) of single-cell RNA-seq data of CD3+CD45+ cells in the blood (**f**) and dissociated thymus (**g**) across all scRNA-seq time points, with annotated cell types. (Bottom) UMAP colored by cell density across representative time points.
**d,** Gene expression of marker genes for annotated cell types in single-cell RNA-seq data of CD45+ sorted peripheral blood.
**e,** Changes in cell types over age in the thymus based on single-cell data, fold changes calculated compared against the youngest time point, 4 weeks.
**f,** Changes in cell type proportions of NKT, CD8+, and CD4+ with age in single-cell sequencing of CD45+ sorted peripheral blood. Fold changes calculated compared against the youngest time point, 4 weeks.
**eg** Changes in cell types over age in the CD45+ sorted peripheral blood based on single-cell data, fold changes calculated compared against the youngest time point, 4 weeks.

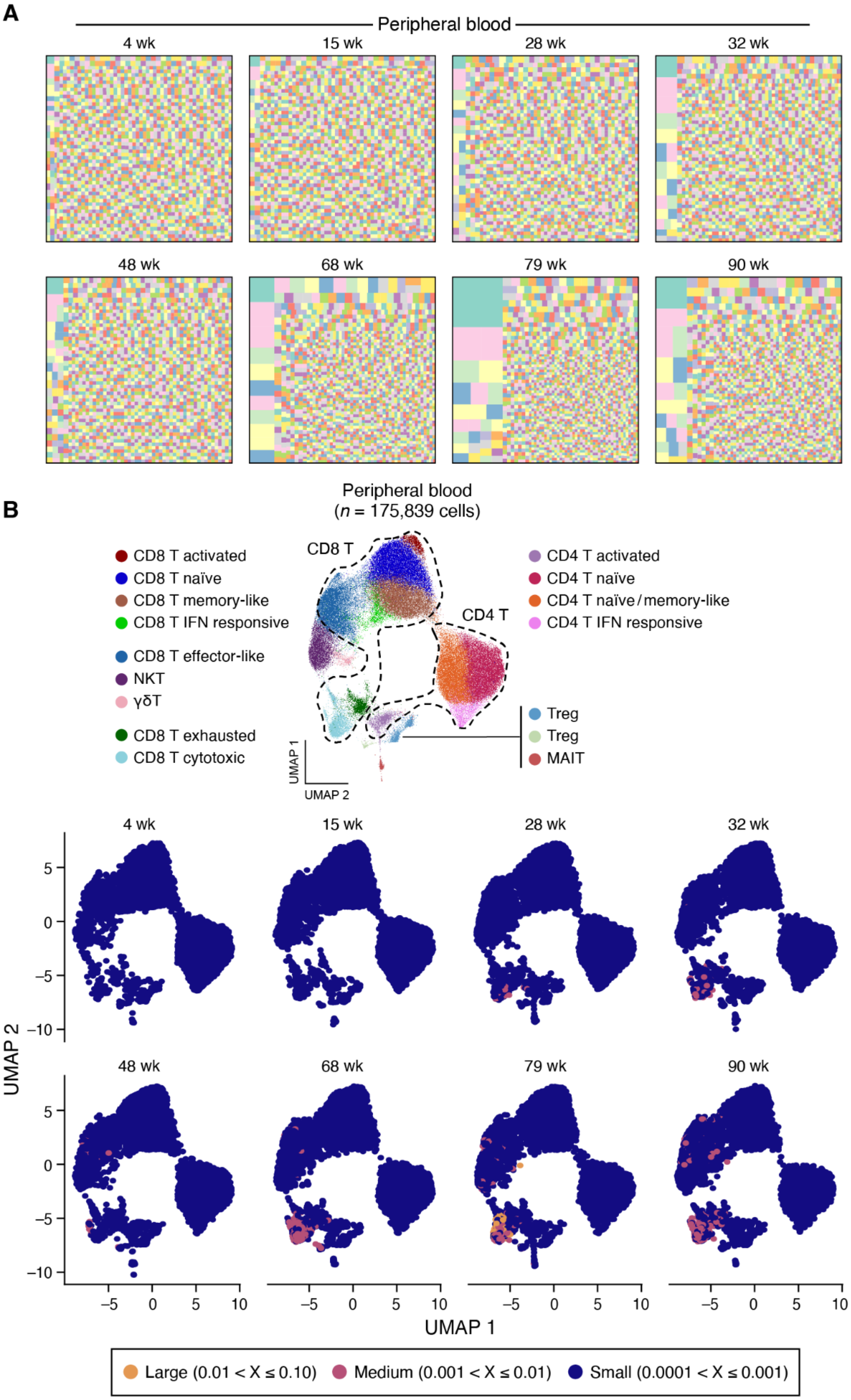
A
Peripheral blood
4 wk
15 wk
28 wk
32 wk
48 wk
68 wk
79 wk
90 wk
B
Peripheral blood
(n = 175,839 cells)
CD8 T
CD4 T
CD8 T activated
CD8 T naïve
CD8 T memory-like
CD8 T IFN responsive
CD8 T effector-like
NKT
γδT
CD8 T exhausted
CD8 T cytotoxic
CD4 T activated
CD4 T naïve
CD4 T naïve/memory-like
CD4 T IFN responsive
Treg
Treg
MAIT
UMAP 1
UMAP 2
UMAP 2
UMAP 1
Large (0.01 < X ≤ 0.10)
Medium (0.001 < X ≤ 0.01)
Small (0.0001 < X ≤ 0.001)

**Extended Data Fig. 7. TCR repertoire shifts in the peripheral blood**

**a,** TCR repertoire diversity in the peripheral blood using CDR3 regions, downsampled to 7500 cells per time point, at representative young (4 weeks) and aged (90 weeks) time points. Each colored square represents a unique clone, with the size of each square representing the fraction of the total repertoire belonging to each clone.
**b,** (Top) UMAP from **Fig. 1f** for reference. (Bottom) UMAP of CD45+ cells from scRNA-seq of peripheral blood over time. Cells are colored by size of each TCR clone.

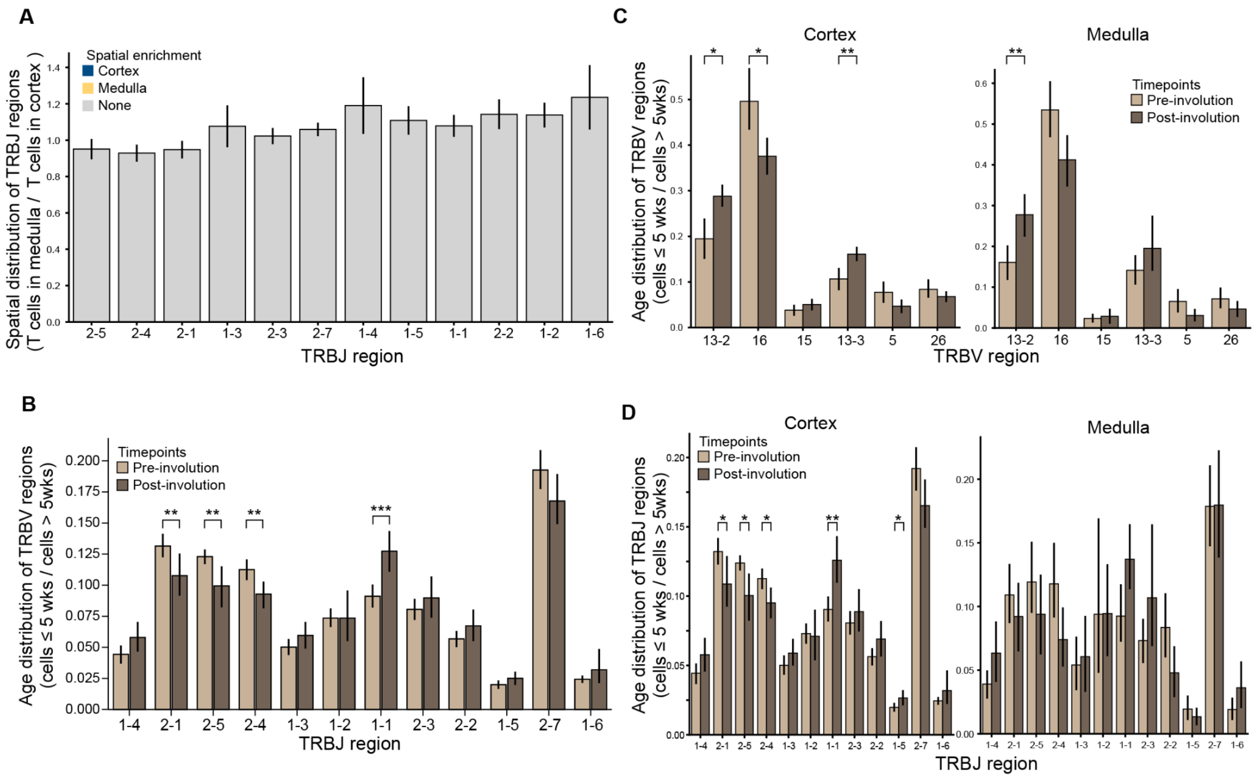


**Extended Data Fig. 8. Spatial alterations in T cell repertoire throughout aging**

**a,** Fraction of TRBJ regions in medullary vs. cortical regions of the thymus across all time points (n=21 mice) in the spatial data. Two-sample paired t test and Benjamini-Hochberg correction used for comparison between TRBV regions. Error bars represent standard error.

**b,** Fraction of TRBJ regions in pre-involution (<= 5 wk, n=12 mice) versus post-involution thymus (>5 wk, n=9 mice). Two-sample t test and Benjamini-Hochberg correction used for comparison between age groups. Error bar represents 95% confidence interval.

**c,** Fraction of TRBV regions in pre-involution (<= 5 wk, n=12 mice) versus post-involution thymus (>5 wk, n=9 mice) separated by cortical (left) and medullary (right) regions. Two-sample t test and Benjamini-Hochberg correction used for comparison between age groups. Error bar represents 95% confidence interval.

**d,** Fraction of TRBJ regions in pre-involution (<= 5 wk, n=12 mice) versus post-involution thymus (>5 wk, n=9 mice) separated by cortical (left) and medullary (right) regions. Two-sample t test and Benjamini-Hochberg correction used for comparison between age groups. Error bar represents 95% confidence interval.

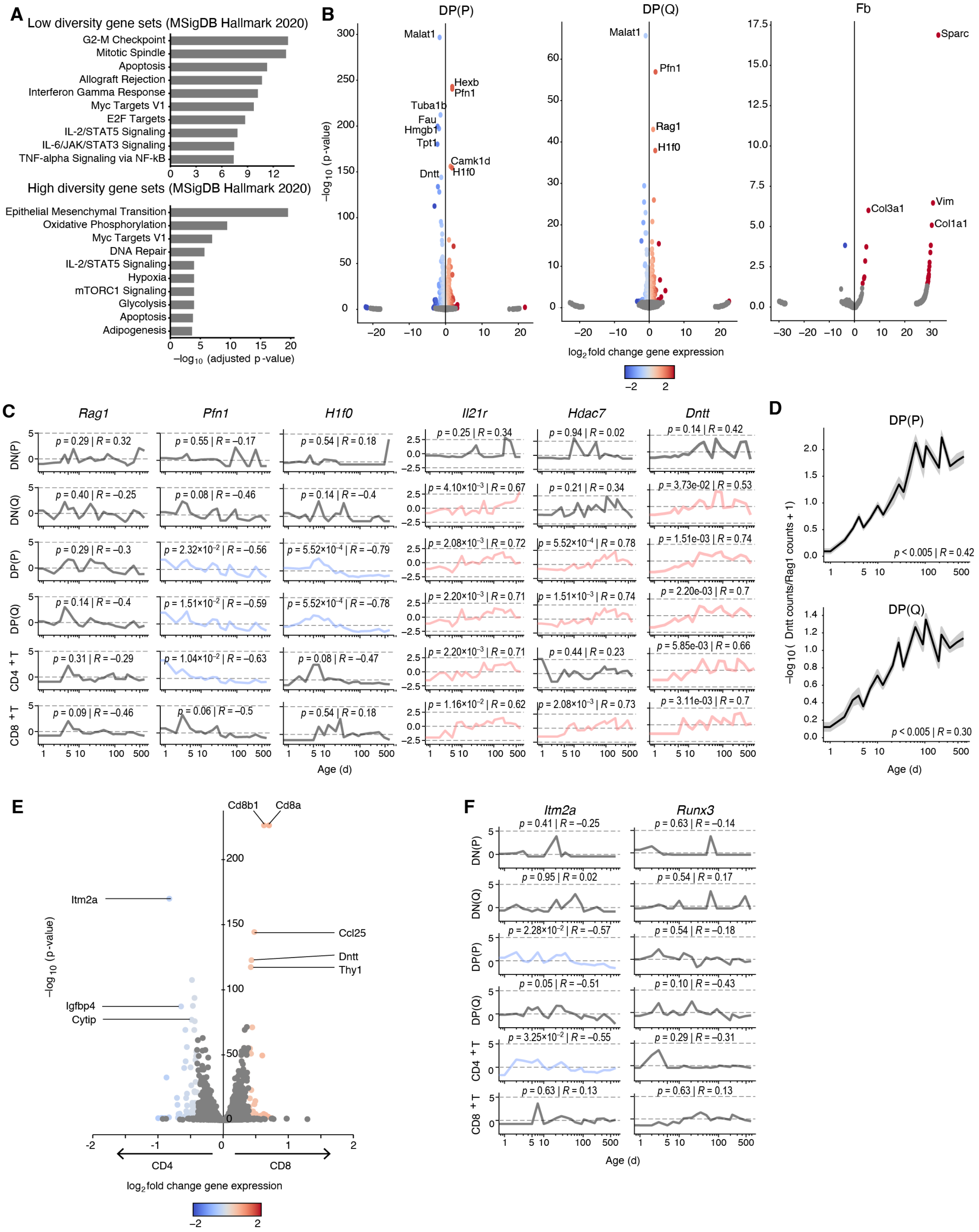


**Extended Data Fig. 9. Changes in thymocyte states with age**

**a,** Gene set enrichment analyses of genes expressed in high and low TCR diversity regions in the thymus.

**b,** Differential gene expression between high (top 10%) and low diversity (bottom 10%) spatial regions of the thymus (n=21 mice), analysis separated by individual cell types. Color bar denotes log2 fold change.
**c,** Changes in gene expression in T cells along the developmental trajectory across age, z-scored across all time points for each gene. p value denotes the significance of Pearson's correlation with respect to age.
**d,** Increase in the expression of *Dntt*, normalized against *Rag1* expression, in DP(P) and DP(Q) cells with age.
**e,** Differential gene expression between CD4+T and CD8+T spatial regions of the thymus.
**f,** Changes in gene expression of Itm2a and Runx3 in T cells along the developmental trajectory across age, z-scored across all time points for each gene. P value denotes the significance of Pearson's correlation with respect to age.
